# Parsing brain-behavior heterogeneity in very preterm born children using integrated similarity networks

**DOI:** 10.1101/2022.10.20.513074

**Authors:** Laila Hadaya, Konstantina Dimitrakopoulou, Lucy Vanes, Dana Kanel, Sunniva Fenn-Moltu, Oliver Gale-Grant, Serena J Counsell, A David Edwards, Mansoor Saqi, Dafnis Batalle, Chiara Nosarti

## Abstract

Very preterm birth (VPT; ≤ 32 weeks’ gestation) is associated with altered brain development and cognitive and behavioral difficulties across the lifespan. However, heterogeneity in outcomes among individuals born VPT makes it challenging to identify those most vulnerable to neurodevelopmental sequelae. Here, we aimed to stratify VPT children into distinct behavioral subgroups and explore between-subgroup differences in neonatal brain structure and function. 198 VPT children (98 females) previously enrolled in the Evaluation of Preterm Imaging study (EudraCT 2009-011602-42) underwent Magnetic Resonance Imaging at term-equivalent age and neuropsychological assessments at 4-7 years. Using an integrative clustering approach, we combined neonatal socio-demographic, clinical factors and childhood socio-emotional and executive function outcomes, to identify distinct subgroups of children based on their similarity profiles in a multidimensional space. We characterized resultant subgroups using domain-specific outcomes (temperament, psychopathology, IQ and cognitively stimulating home environment) and explored between-subgroup differences in neonatal brain volumes (voxel-wise Tensor-Based-Morphometry), functional connectivity (voxel-wise degree centrality) and structural connectivity (Tract-Based-Spatial-Statistics). Results showed two-and three-cluster data-driven solutions. The two-cluster solution comprised a ‘resilient’ subgroup (lower psychopathology and higher IQ, executive function and socio-emotional outcomes) and an ‘at-risk’ subgroup (poorer behavioral and cognitive outcomes). The three-cluster solution showed an additional third ‘intermediate’ subgroup displaying behavioral and cognitive outcomes intermediate between the resilient and at-risk subgroups. The resilient subgroup had the most cognitively stimulating home environment and the at-risk subgroup showed the highest neonatal clinical risk, while the intermediate subgroup showed the lowest clinical but the highest socio-demographic risk. Compared to the intermediate subgroup, the resilient subgroup displayed larger neonatal insular and orbitofrontal volumes and stronger orbitofrontal functional connectivity, while the at-risk group showed widespread white matter microstructural alterations. These findings suggest that risk stratification following VPT birth is feasible and could be used translationally to guide personalized interventions aimed at promoting children’s resilience.

## 1. Introduction

Very preterm birth (VPT; ≤ 32 weeks’ gestation) is associated with an increased likelihood of developing cognitive and behavioral difficulties across the lifespan (^1–5^). Efforts to conceptualize these difficulties have proposed a “preterm behavioral phenotype”, characterized by deficits in emotional and social processing, and inattention (^6^). However, while some VPT children display a behavioral profile reflecting a preterm phenotype, others follow typical developmental trajectories (^7–9^). Such behavioral heterogeneity following VPT birth presents a challenge for building risk prediction models (^10^), as multiple causes may lead to the same outcome and as a single mechanism may lead to multiple outcomes (^11^).

Several endogenous and exogenous factors contribute to a child’s behavioral development and a complex interplay between environmental, clinical, and neurobiological features could result in co-occurring neurodevelopmental, cognitive and behavioral difficulties following VPT birth (^12^). These factors are often non-independent and their combination (e.g., neurobiological and socio-demographic variables) may result in improved prediction of functional outcomes (^13^). For instance, both socio-demographic deprivation and increased neonatal clinical risk have been associated with neurodevelopmental as well as behavioral difficulties in VPT children. These encompass executive and socio-emotional functions (^14–16^), which could be considered as gateway mechanisms that shape behavioral outcomes, as they are subserved by brain networks relating to both bottom-up stimulus processing and top-down behavioral control (^17^). Impairments in these domains have in fact been associated with later mental health and academic difficulties (^3,18^).

Previous studies have attempted to stratify heterogeneity amongst preterm children using clustering and latent-class analyses, based on socio-demographic, clinical, socio-emotional, and cognitive measures (^7–9,19,20^). While some studies found differences in neonatal clinical profiles between subgroups of VPT children (^20^), others showed that familial characteristics such as parental education, maternal distress, and cognitively stimulating parenting differentiated resilient subgroups from those exhibiting behavioral difficulties (^8,9^). However, while socio-demographic and/or environmental factors may explain a proportion of functional outcomes, a growing body of research is studying the early neural signatures that may shape an individual’s neurodevelopmental trajectory. Alterations in brain volumes (^21,22^), white matter microstructure (^23,24^), and functional connectivity (^25,26^) at birth in regions and networks subserving social, emotional and attentional processes, have been associated with later behavioral difficulties in VPT samples. However, it remains to be explored whether distinct multidimensional subgroups of VPT children could also be characterized by differences in early brain development.

Therefore, this study set out to parse brain-behavior heterogeneity in VPT children. Firstly, we implemented an integrative clustering approach (Similarity Network Fusion; SNF) (^27^) to stratify VPT children into distinct subgroups based on three data types: i) neonatal clinical and socio-demographic variables, ii) childhood socio-emotional outcomes and iii) executive function measures. The advantage of this approach is that it integrates sample-similarity networks built from each distinct data type and constructs a final integrated network, which contains common and complementary information from the different data types. This is then used to stratify the sample into distinct subgroups using clustering (^27^). We also investigated whether resultant subgroups differed in outcomes that were not used in stratification analyses (i.e., out-of-model variables); in order to provide external validation (^28–30^). Finally, we explored between-subgroup differences in regional brain volume and structural and functional connectivity at term-equivalent age. We hypothesized that there would be distinct subgroups of VPT children characterized by unique neonatal neural signatures.

## 2. Methods

### 2.1 Study design

#### 2.1.1 Participants

511 infants born VPT were recruited from 14 neonatal units in London in 2010-2013 and entered the Evaluation of Preterm Imaging Study (ePrime; EudraCT 2009-011602-42) (^31^). Infants with congenital malformation, prior Magnetic Resonance Imaging (MRI), metallic implants, whose parents did not speak English or were subject to child protection proceedings were not eligible for participation in the study.

Participants underwent multimodal MRI scanning at 38-53 weeks post-menstrual age (PMA) on a 3-Tesla MR imaging system (Philips Medical Systems, Best, The Netherlands) located on the neonatal intensive care unit at Queen Charlotte’s and Chelsea Hospital, London, using an 8-channel phased array head coil. For data acquisition and imaging parameters see Supplemental Information. Infants whose parents chose sedation for the procedure (87%) received oral chloral hydrate (25–50 mg/kg).

251 participants (including 29 sets of multiple pregnancy children) were followed-up between the age of 4 and 7 years at the Centre for the Developing Brain, St Thomas’ Hospital, London. This was a convenience sample corresponding to 82% of 306 participants who were eligible and invited to participate.

Ethical approval was granted by the Hammersmith and Queen Charlotte’s Research Ethic Committee (09/H0707/98) and the National Research Ethics Committee (14/LO/0677).

#### 2.1.2 Clinical and socio-demographic data

The following neonatal clinical variables were evaluated: gestational age (GA) at birth, number of days on mechanical ventilation, number of days on continuous positive airway pressure (CPAP) and number of days on parenteral nutrition (TPN). Please refer to Supplemental Information for more details.

Socio-demographic risk was evaluated using a postcode derived measure of deprivation in England, the Index of Multiple Deprivation 2010 (IMD; http://tools.npeu.ox.ac.uk/imd/), whereby higher IMD scores reflect greater deprivation. The IMD combines neighbourhood-specific information about seven domains of deprivation: income, employment, education/skills/training, health, crime, housing and living environment. Continuous IMD scores were used in the integrative-clustering and evaluation of subgroup profile analyses. IMD quintiles are provided when reporting sample characteristics (Table 1) for ease of interpretability.

**Table 1.**
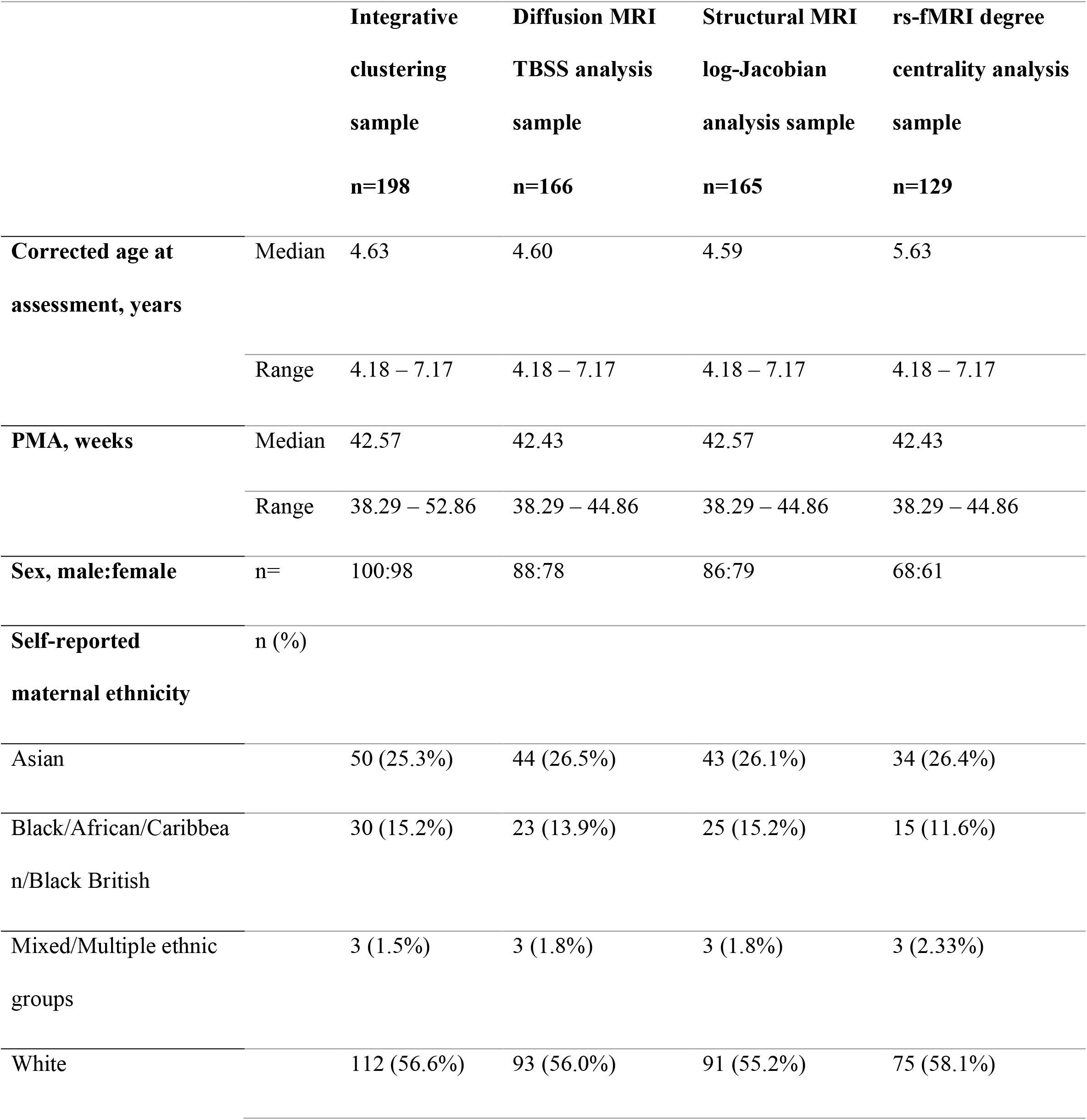
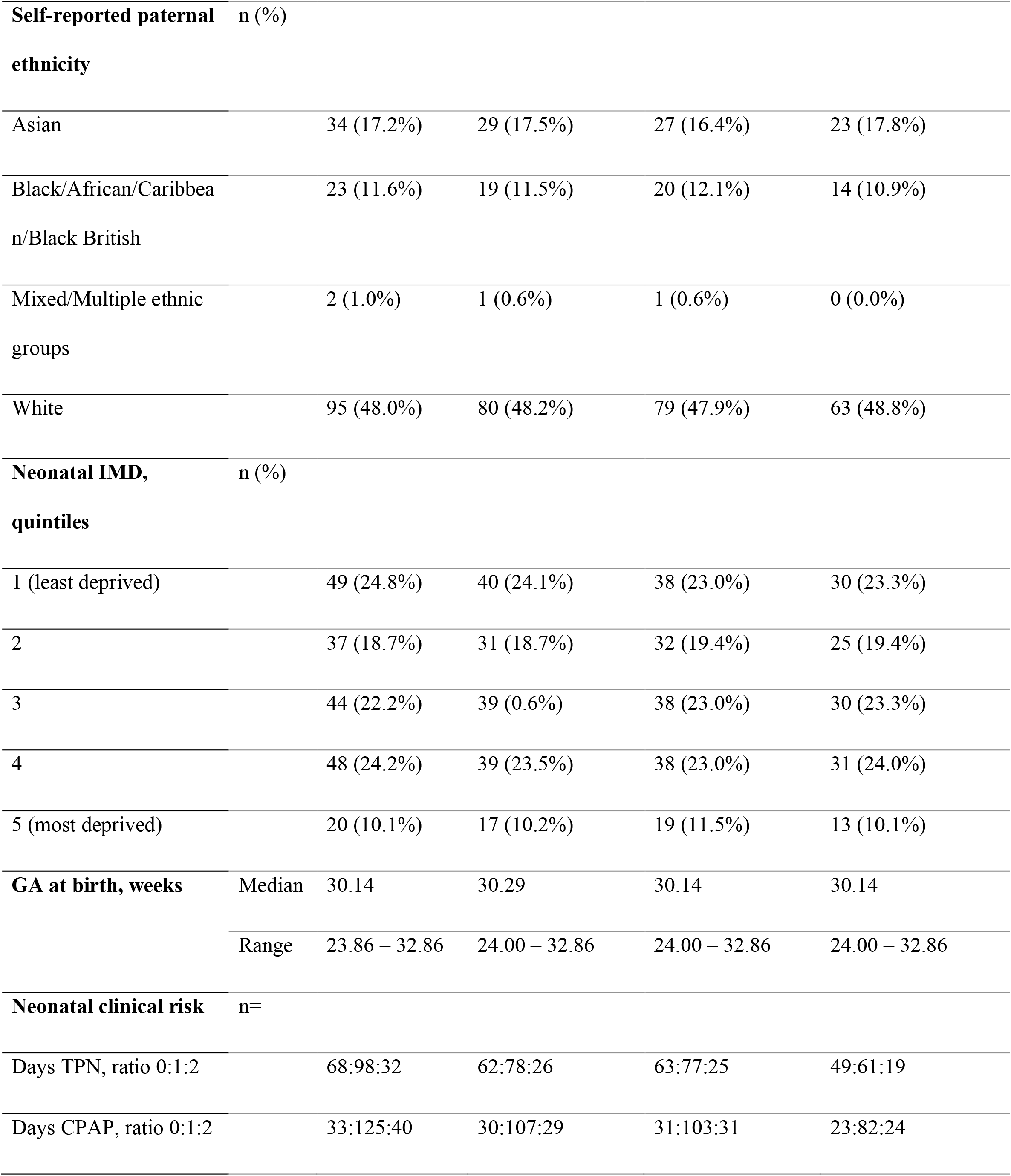
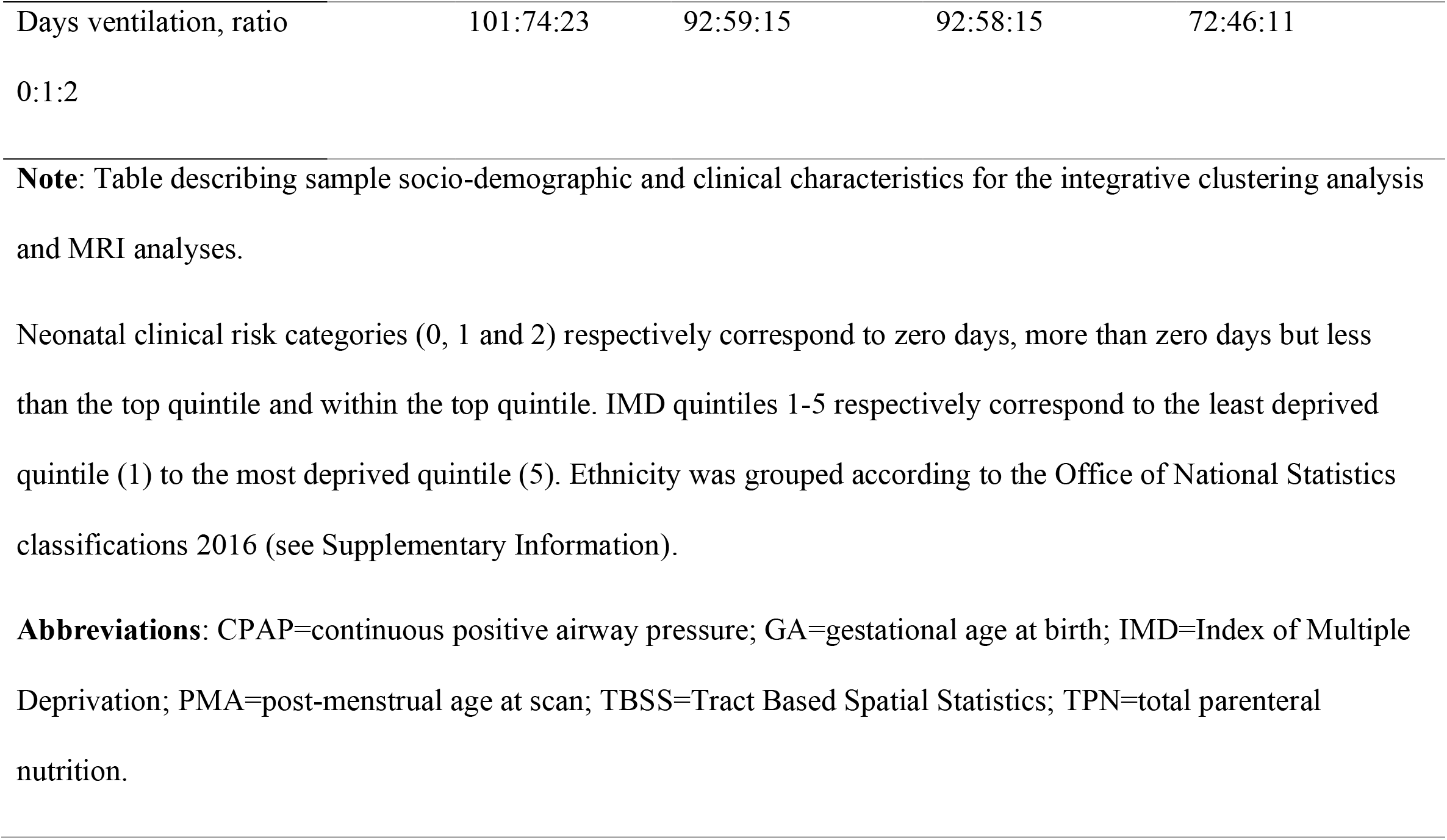
Socio-demographic and clinical participant data.

#### 2.1.3 Childhood assessment

Intelligence quotient (IQ) was assessed using the Wechsler Preschool and Primary Scale for Intelligence (WPPSI-IV) (^32^) and executive function using the preschool version of the parent-rated Behaviour Rating Inventory of Executive Function (BRIEF-P) (^33^). Socio-emotional processing was evaluated using the Empathy Questionnaire (EmQue) (^34^) and the Social Responsiveness Scale, Second Edition (SRS-2) (^35^). Psychopathology was assessed using the Strengths and Difficulties Questionnaire (SDQ) (^36^), temperament using the Child Behavioural Questionnaire - Very Short Form (CBQ) (^37^) and cognitively stimulating home environment using an adapted version of the Cognitive Stimulating Parenting Scale (CSPS) (^38^).

#### 2.1.4 Exclusions

Twenty-seven participants were excluded due to incomplete childhood outcome data, 17 due to major brain lesions (periventricular leukomalacia, parenchymal hemorrhagic infarction, or other ischemic or hemorrhagic lesions) detected on neonatal T2-weighted MRI images at term by an experienced perinatal neuroradiologist, and 5 participants due to missing T2-weighted MRI images, hence inability to evaluate the presence of major lesions (Figure S1).

### 2.2 Data integration and clustering

Analyses were conducted in R (version 3.6.1). Using SNF, three data types were integrated: **Type 1)** neonatal socio-demographic and clinical variables: IMD at birth, GA, days on ventilation, days on TPN and days on CPAP. **Type 2)** childhood socio-emotional outcomes: EmQue subscale raw scores - emotion contagion, attention to others’ emotions, prosocial behaviors and SRS-2 total raw score. **Type 3)** childhood executive function: BRIEF raw subscale scores - inhibit, shift, emotional control, working memory and plan/organize.

Prior to integration, participants with in-model outlier values greater than 3 times the interquartile range were excluded. A total of 198 children were included in the SNF analyses. Zero-inflated neonatal clinical risk variables (days ventilation, days TPN and days CPAP) were converted into ordinal categorical variables with three levels: (Level 0: zero days; Level 1: greater than zero and not within top quintile; Level 2: within top quintile). For the mixed data type (numeric and categorical data; data type 1), Gower’s standardization based on the range was applied using the *daisy* function from cluster R package (^39^) and for numeric only matrices (data types 2 and 3), variables were standardized to have a mean value of 0 and a standard deviation of 1 using the *standardNormalization* function from SNFtool R package (^40^).

An adaptation of the *ExecuteSNF*.*CC* function (^41^) was used for the data integration and clustering steps. Dissimilarity Gower distance (for the mixed data type) and Euclidean distance (for numeric data types) matrices were calculated and used to create similarity matrices using the SNFtool R package’s *affinityMatrix* function (^40^). This was followed by an integration of the similarity matrices using SNFtool’s *SNF* algorithm resulting in a ‘fused similarity matrix’ (^40^). The integrative clustering process can be summarized into two steps:

**Step 1**: SNF method has two main hyperparameters, K and alpha. K (i.e., neighborhood size) indicates the number of neighbors of a node to consider when the similarity networks are being generated and alpha is an edge weighting parameter determining the weight of edges between nodes in the networks. We tried 30 combinations of K and alpha hyperparameters {K=10, 15, 20, 25, 30; alpha=0.3, 0.4, 0.5, 0.6, 0.7, 0.8}, similar to the approach followed in (^42^). The K-alpha hyperparameter values were chosen based on the ranges recommended in the SNFtool R package, 10-30 for K and 0.3-0.8 for alpha (^27,40^). Consensus clustering, using *ConsensusClusterPlus* function (^43^), was then applied to each fused similarity matrix, corresponding to a K-alpha combination, where spectral clustering was run 1000 times with 80% of the population randomly subsampled for each clustering run and a single consensus clustering result obtained from hierarchical clustering. **Step 2**: Next, out of the 30 clustering results produced in step 1, the one with the highest average silhouette width score was retained. Steps 1 and 2 were repeated 1000 times in a bootstrap approach, after selecting and pre-processing the three data matrices of 80% of the sample set. The 1000 resultant retained clustering outputs were then fed to the diceR R package’s *consensus_combine* function (^44^) which implements hierarchical clustering on the consensus matrix and generates the final consensus clustering. Figure 1 summarizes the data-integration and clustering steps and the code used can be accessed here: https://github.com/lailahadaya/preterm-ExecuteSNF.CC.xs Further details can also be found in the Supplemental Information.

**Figure 1.**
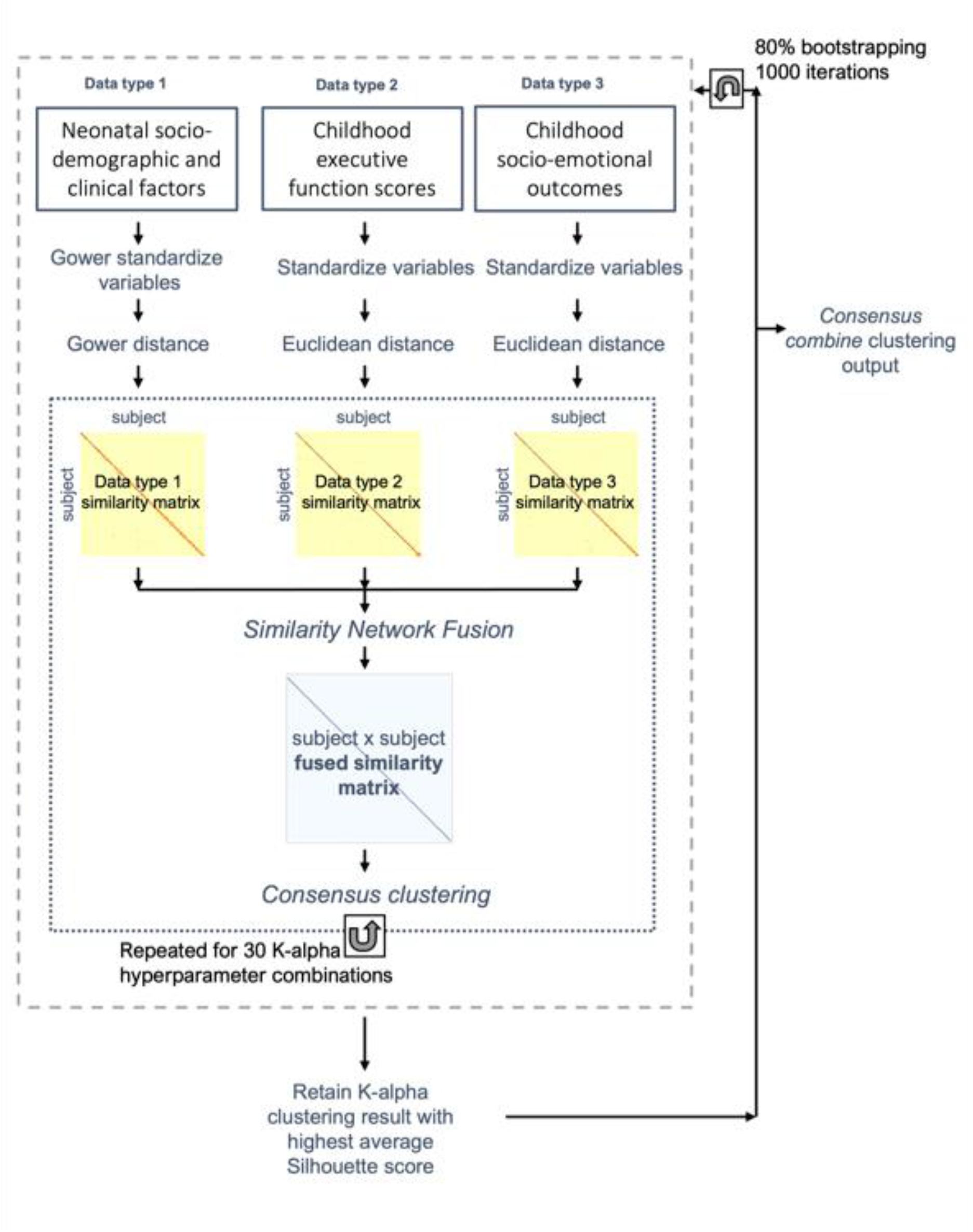
Data integration and clustering pipeline. Figure summarizing the data pre-processing (variable normalization), data integration and clustering pipeline executed in order to obtain the final consensus cluster assignment.

Before implementing steps 1 and 2, it was essential to determine the number of clusters. For this, we used the SNFtool R package’s *estimateNumberOfClustersGivenGraph* function (^40^) to calculate Eigengap and Rotation Cost heuristics for each K-alpha combination (Figure S2). This process suggested C=2, C=3 and C=4 as the optimal number of clusters. Consensus matrices and silhouette scores were generated and compared for these three potential clustering solutions (Figure S2). Resultant subgroups from C=2 and C=3 were chosen to be evaluated for phenotypic differences, as their silhouette scores and consensus matrices gave better values in comparison to those of C=4 (Figure S2). More details on the estimation of cluster numbers can be found in the Supplemental Information. An alluvial plot was used to illustrate the transition of subject subgroup classification between the two-cluster and three-cluster solutions (Figure S3).

### 2.3 Evaluation of subgroup profiles

Resultant subgroups were characterized based on in-model and out-of-model variables. For the out-of-model features, subgroups were compared in terms of psychiatric symptoms (SDQ internalizing, externalizing problems and total scores), temperament (CBQ negative affectivity, surgency and effortful control scores), cognitive abilities (WPPSI full-scale IQ), and cognitive stimulation at home (CSPS score). Details on selection of in-model and out-of-model variables can be found in Supplemental Information and Figures S4 and S5.

For numeric measures, between-subgroup differences were assessed using non-parametric one-way tests: Mann-Whitney when C=2 or Kruskal Wallis when C=3 (^45^). Shapiro-Wilk test was used to assess normality. For categorical variables, Chi-squared test was used to evaluate differences in proportions of individuals in each group when count per cell was >5 and Fischer’s Exact test was used otherwise. To compare differences between the ordinal neonatal clinical variables with 3 categories (Levels 0, 1 and 2) and the non-ordinal subgroups from C=2 and C=3, the Extended Cochran-Armitage Test was used.

Results with *p*<0.05 were considered to be statistically significant. To correct for multiple comparisons the False Discovery Rate method was used. The same statistical analyses were repeated using general linear models correcting for potential confounders (age and sex) and 5000 permutation test iterations (^46^). Effect sizes for non-normally distributed variables were measured using Wilcoxon Glass Rank Biserial Correlation (gr) for measuring differences between two groups and Epsilon Squared for three groups. For normally distributed variables, Cohen’s F was used and Cramer’s V for categorical variables.

### 2.4 Exploring neonatal brain differences between subgroups

Tract Based Spatial Statistics (TBSS) was used to assess white matter microstructure at the voxel-level using fractional anisotropy (FA) and mean diffusivity (MD) maps (^47^). FA approximates the directional profile of water diffusion in each voxel and MD measures the average movement of water molecules within a voxel. Higher FA and lower MD values reflect more optimal white matter myelination and microstructure. For diffusion MRI (d-MRI) image pre-processing and TBSS protocol details please refer to Supplemental Information.

Structural MRI (s-MRI) log-Jacobian determinant maps were calculated to quantify regional brain volumes (greater log-Jacobian values reflect larger relative structural volumes), using Tensor Based Morphometry, following methods described in our previous work (^48,49^) and in Supplemental Information.

Resting-state functional MRI (rs-fMRI) data were pre-processed as in our previous work (^50^); for more details see Supplemental Information. Functional connectivity was evaluated using a measure of weighted degree centrality at the voxel-level (i.e., the sum of the correlations between the time-series of each voxel and all other voxels within a grey matter mask of the brain) (^51,52^). Edges with a correlation coefficient below a threshold of 0.2 were excluded and the degree centrality values for each voxel in the grey matter mask were z-scored and used in subsequent between-subgroup analyses.

The number of children included in the different modality-specific MRI analyses slightly differed due data availability: TBSS (n=166), log-Jacobian determinant maps (n=165) and degree centrality (n=129); please see Table S1. Exclusions for specific MRI analyses are depicted in Figure S1.

Between-subgroup differences were investigated in the whole-brain at the voxel-level in terms of: log-Jacobian determinants, TBSS metrics (FA and MD) and degree centrality. FMRIB Software Library (FSL) (^53^) *randomise* function was used to implement non-parametric permutation methods for statistical inference. This method was used to model each contrast of interest for each voxel, i.e., a general linear model (GLM) correcting for PMA at scan and sex. rs-fMRI models also included motion estimates (standardized DVARS) as a covariate. Family Wise Error (FWE) rate with Threshold-Free Cluster Enhancement was applied to correct for multiple comparisons over the multiple voxels, while enhancing “cluster-like” structures of voxels without defining them as binary components (^54^). Statistics were calculated using random permutation tests with 10000 permutations. Given the exploratory nature of our analysis, we did not correct for multiple contrasts tested (i.e., log-Jacobians, TBSS FA and MD, degree centrality). We show results significant at *p*<0.05 FWE-corrected per contrast. Mean values from clusters of modality-specific voxels showing significant between-subgroup differences were extracted to calculate Cohen’s F effect sizes.

### 2.5 Sensitivity analyses

There were 29 sets of children born from multiple pregnancy events in our sample. In order to account for multiple pregnancy confounding, we conducted additional sensitivity analyses including only one child from each set of multiple pregnancy siblings.

## 3. Results

### 3.1 Participant characteristics

Participants’ socio-demographic and clinical characteristics are shown in Table 1. Compared to those who returned for follow-up (n=251; median GA=29.24 weeks; median IMD at birth=19.48), non-returners (n=259; median GA=29.27 weeks; median IMD at birth=21.40) did not differ in GA (gr=0.01; *p*=0.807), but had greater neonatal socio-demographic deprivation (gr=0.11; *p*=0.028). Compared to the initial baseline cohort (n=511; median GA=30.00 weeks; median IMD at birth=18.19), participants who were studied here (n=198) had slightly older GA (median GA=30.14 weeks; gr=-0.13; *p*=0.009) and relative socio-demographic advantage (median IMD score at birth=15.58, *gr*=0.11; *p*=0.027).

### 3.2 Two-cluster solution subgroup profiles

When stratifying the sample into two clusters and comparing them in terms of in-model variables, subgroup 1 (termed here the ‘resilient’ subgroup) showed significantly better socio-communication (i.e., lower SRS-2 scores) and executive function abilities (i.e., lower BRIEF emotion control, inhibit, shift, working memory and plan/organize scores), lower emotion contagion (EmQue) scores, and higher prosocial actions scores (EmQue) during childhood, than subgroup 2 (termed here the ‘at-risk’ subgroup); all *p*s<0.05, even after FDR correction. The resilient subgroup had lower neonatal clinical risk compared to the at-risk subgroup, with a greater proportion of children receiving no neonatal mechanical ventilation and a smaller proportion of children receiving prolonged neonatal CPAP (both *ps*<0.05, even after FDR correction). Subgroups did not differ in terms of days on TPN in the neonatal period (*p*>0.05).

Differences in out-of-model variables included lower psychopathology scores (SDQ internalizing and externalizing problems) and negative affectivity scores (CBQ) as well as higher effortful control (CBQ), IQ and cognitive stimulation at home (CSPS) during childhood in the resilient compared to the at-risk subgroup; all *p*s<0.05, even after FDR correction (Figure 2; Table S2).

**Figure 2.**
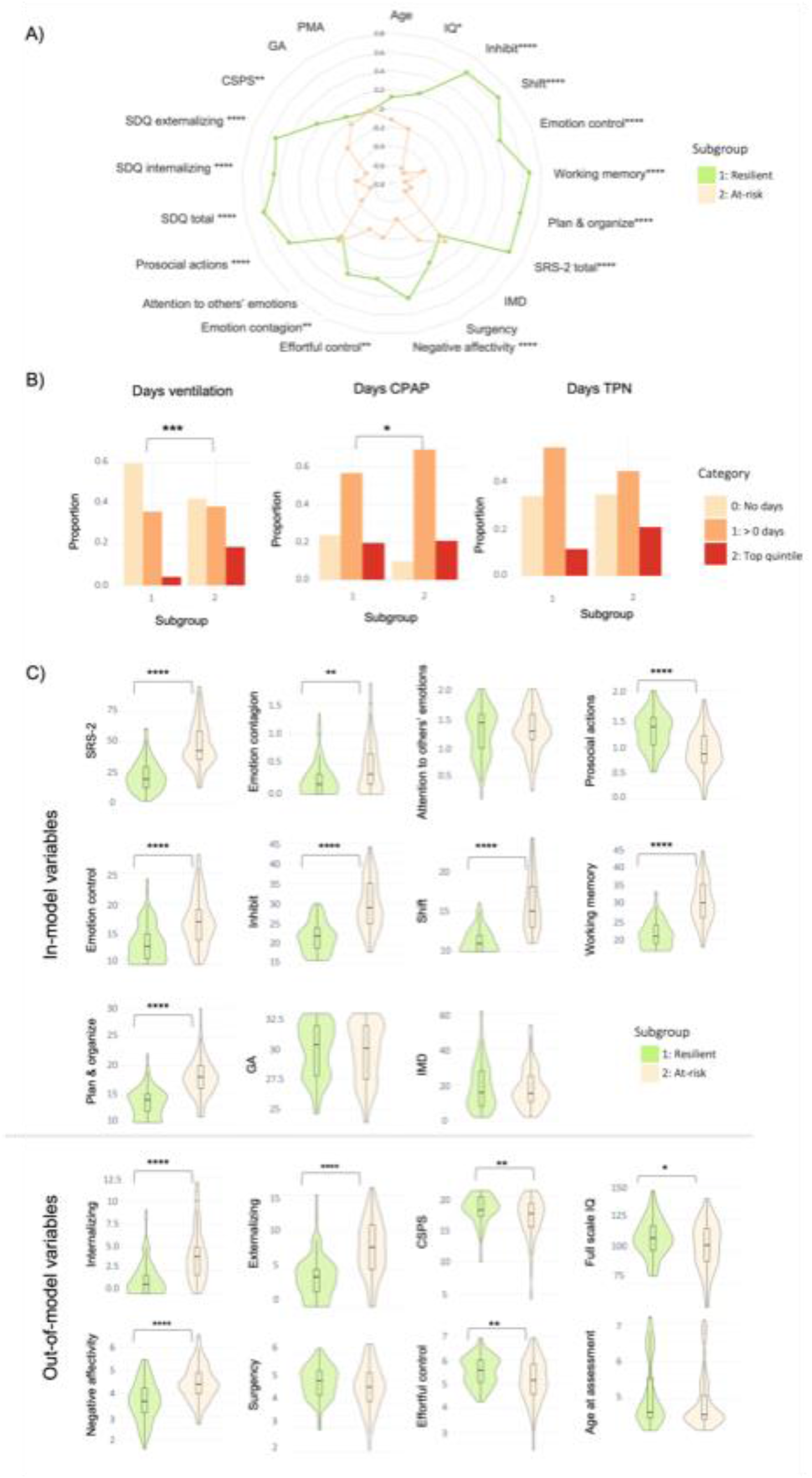
Two-cluster solution subgroup profiles. A) Radar plot showing the two-cluster solution subgroup profiles using z-scores for subgroup 1 (i.e., resilient subgroup; green) and subgroup 2 (i.e., at-risk subgroup; beige). For visual illustrative purposes, scales which usually indicate poorer outcomes have been reversed so that larger z-scores on behavioral variables indicate better outcomes. B) Bar plots for clinical risk variables (days on TPN, days on mechanical ventilation and days on CPAP, left to right, respectively) for each of the two subgroups. Plots represent the proportion of children belonging to each clinical risk category within a subgroup, where category 0 represents the lowest clinical risk (light beige; no days of clinical intervention), category 1 represents medium clinical risk (orange; more than 0 days of intervention but less than the top quintile), and category 2 represents the highest clinical risk (red; within the top quintile). C) Violin plots showing differences between the subgroups in terms of in-model and out-of-model variables. Significant differences are marked with bars between the subgroups. *=*p*<0.05; **=*p*<0.01; ***=*p*<0.001, ****=*p*<0.0001.

The two subgroups showed no significant differences in log-Jacobian determinant values, degree centrality or white matter microstructural characteristics (all *p*s>0.05). Resultant subgroups also did not show differences in sex, age at assessment or PMA at scan (Figure 2; Table S2).

### 3.3 Three-cluster solution subgroup profiles

To increase subtyping resolution and explore latent heterogeneity not captured by a two-subgroup partitioning, the sample was further stratified into 3 subgroups. Two of the three resulting clusters largely reflected profiles similar to those from C=2. The first was a ‘resilient’ subgroup (subgroup 1) with favorable childhood socio-communicative (SRS-2), empathy (EmQue) and executive function (BRIEF) outcomes in terms of in-model variables; low childhood psychopathology (SDQ internalizing and externalizing problems) and negative affectivity scores (CBQ) and high effortful control scores (CBQ), IQ and cognitive stimulation at home (CPSP) in terms of out-of-model variables. The second was an ‘at-risk’ subgroup (subgroup 2), with the poorest outcomes in terms of in-model variables (childhood socio-communication (SRS-2), empathy and executive function (BRIEF) scores), as well as out-of-model childhood psychopathology (SDQ), effortful control (CBQ) and negative affectivity measures (CBQ), combined with the highest neonatal clinical risk (Figure 3; Table S3).

**Figure 3.**
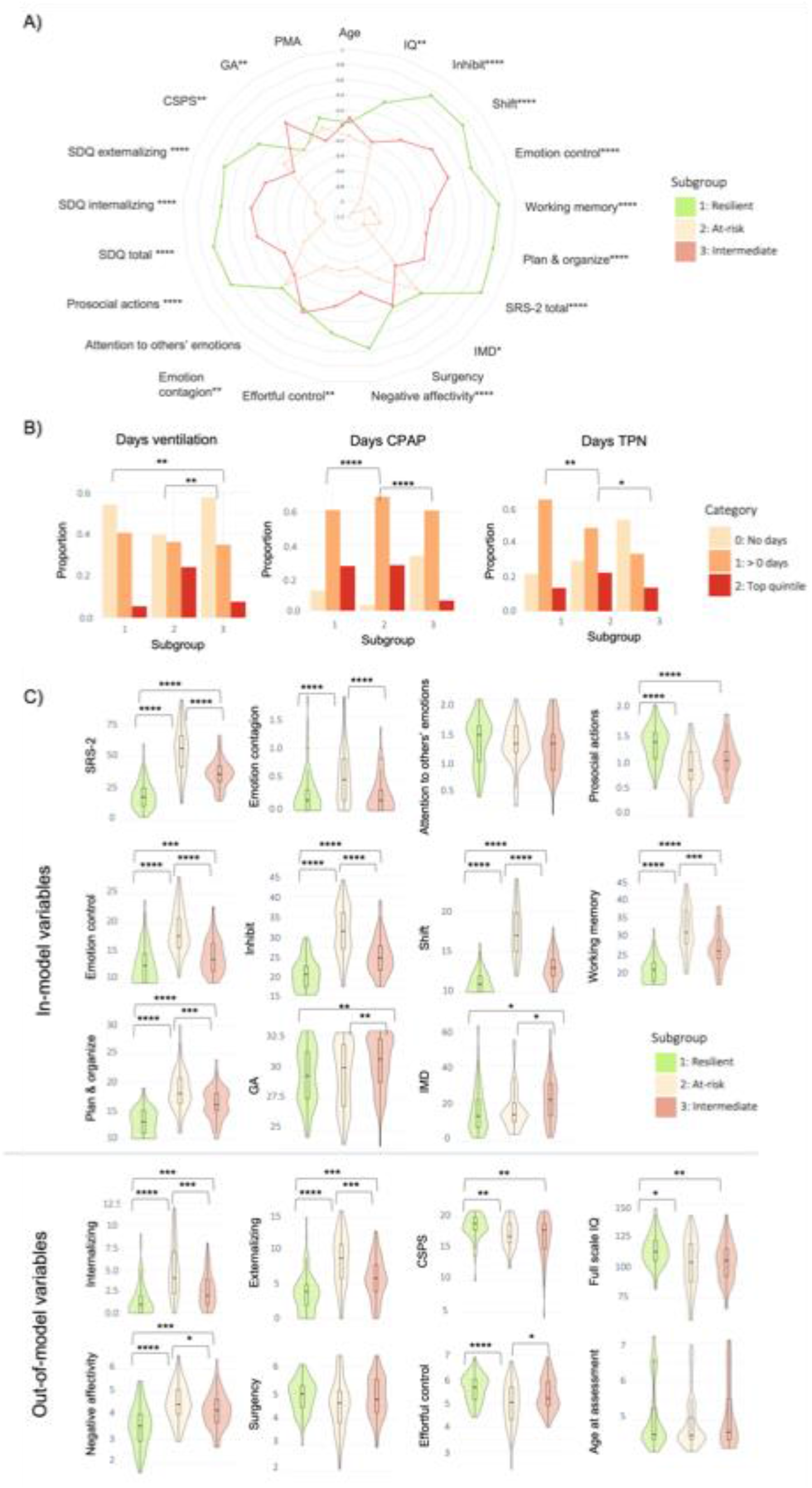
Three-cluster solution subgroup profiles. A) Radar plot showing the three-cluster solution subgroup profiles using z-scores. For visual illustration purposes, scales which usually indicate poorer outcomes have been reversed, so that larger z-scores on behavioral variables indicate better outcomes. Subgroup 1 (resilient) is marked in green, subgroup 2 (at-risk) in beige and subgroup 3 (intermediate outcomes but lowest clinical risk) in pink. B) Bar plots for clinical risk variables (days on TPN, days on mechanical ventilation and days on CPAP, left to right, respectively) for each of the three subgroups. Plots represent the proportion of children belonging to each clinical risk category within a subgroup, where category 0 represents the lowest clinical risk (light beige; no days of clinical intervention), category 1 represents medium clinical risk (orange; more than 0 days of intervention but less than the top quintile), and category 2 represents the highest clinical risk (red; within the top quintile). C) Violin plots present the differences in in-model and out-of-model measures at the group-wise level. Significant differences are marked with bars between the subgroups. *=*p*< 0.05; **=*p*<0.01; ***=*p*<0.001, ****=*p*<0.0001.

A third subgroup (labelled ‘intermediate’) emerged, which had poorer in-model and out-of-model childhood cognitive and behavioral scores when compared to the resilient subgroup, but better scores when compared to those of the at-risk subgroup. The intermediate subgroup also had the lowest neonatal clinical risk compared to both resilient and at-risk subgroups (Figure 3; Table S3). The transition of subject classifications from the two-to the three-cluster solution is illustrated in an alluvial plot (Figure S3).

In terms of environmental factors, the resilient subgroup had higher levels of childhood cognitive stimulation at home (CSPS) in comparison to both at-risk and intermediate subgroups, while the intermediate subgroup had higher neonatal socio-demographic risk (IMD) in comparison to both at-risk and resilient subgroups. The three subgroups did not differ in sex, age at assessment or PMA at scan. All *p*s<0.05 after FDR correction.

In terms of brain imaging markers at term, the resilient subgroup displayed larger relative volumes (i.e., greater log-Jacobian determinant values) in the left insula and bilateral orbitofrontal cortices (Figure 4A; Table S4) and higher degree centrality in an overlapping region in the left orbitofrontal cortex (Figure 4B; Table S5) compared to the intermediate subgroup. The intermediate subgroup, compared to the at-risk subgroup, showed increased FA in several areas of the white matter skeleton, including the fornix, corpus callosum, corticospinal tract, inferior longitudinal, inferior fronto-occipital and uncinate fasciculi (Figure 4Ci; Table S4), as well as lower MD in the fornix and body of the corpus callosum (Figure 4Cii; Table S4). The resilient and at-risk subgroups did not differ in any brain measures (*p*>0.05).

**Figure 4.**
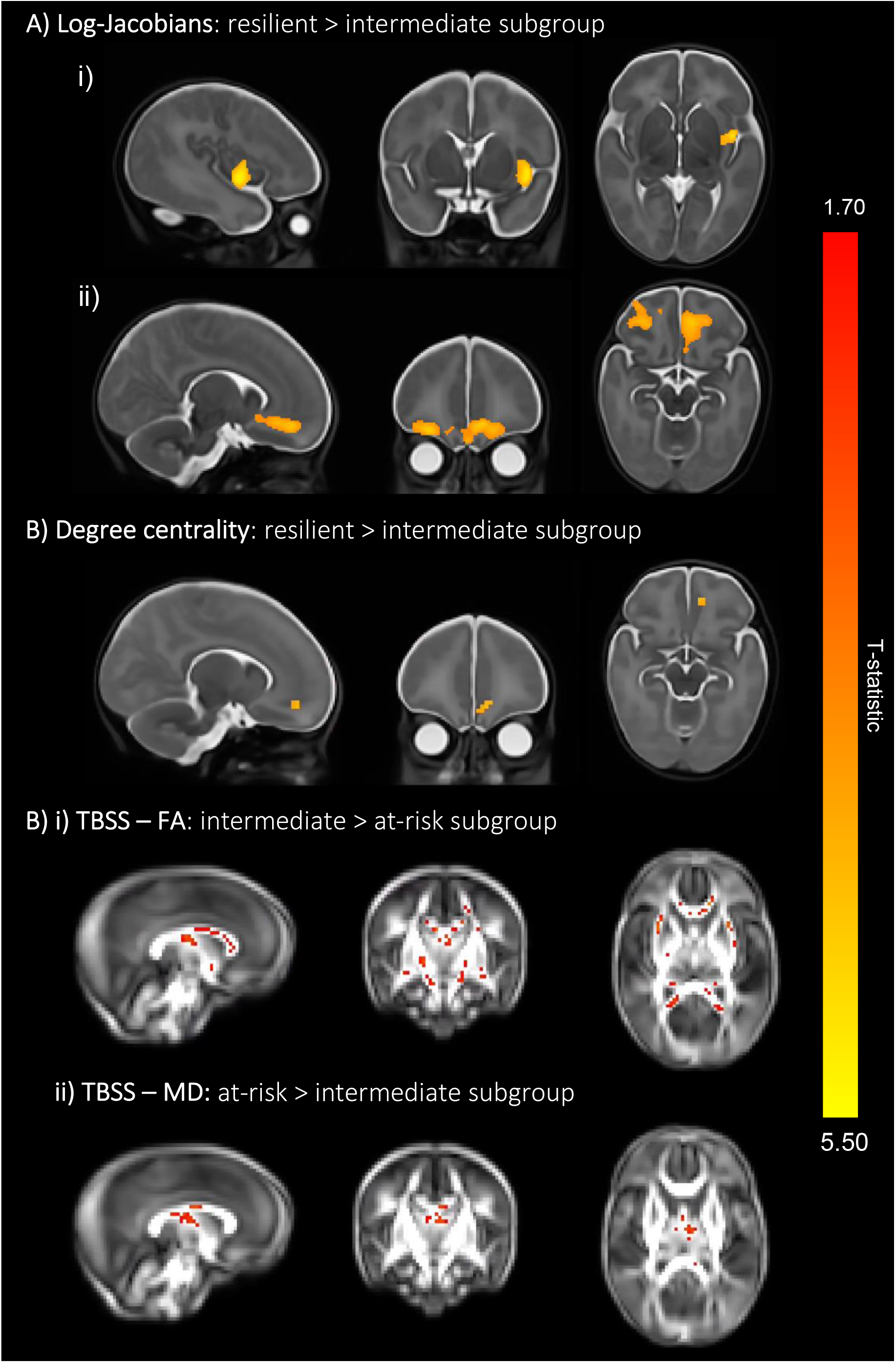
Three-cluster solution brain differences at term-equivalent age. A) Colored voxels indicate regions with significantly larger log-Jacobian determinant values in the resilient subgroup (subgroup 1) compared to the intermediate subgroup (subgroup 3) in i) left insula and the ii) bilateral orbitofrontal cortices (*p*<0.05). GLM included sex and PMA at scan as covariates and TFCE and FWE corrections were applied. B) Voxels showing significantly larger degree centrality values in the resilient subgroup (subgroup) 1 compared to the intermediate subgroup (subgroup 3) are seen in an overlapping left orbitofrontal region at *p*<0.05. GLM included sex, PMA at scan and motion (standardized DVARS) as covariates; TFCE and FWE were applied. C) Colored voxels represent white matter regions showing i) significantly higher FA values in the intermediate subgroup compared to the at-risk subgroup and ii) significantly higher MD values in the at-risk subgroup compared to the intermediate subgroup (*p*<0.05). T-statistic values are represented in the color bar, where red colored voxels indicate smaller T-statistic values and yellow voxels indicate higher T-statistic values ranging between 1.70 and 5.50.

### 3.4 Sensitivity analyses

Sensitivity analyses including only one sibling, selected at random from each multiple pregnancy set, revealed similar results (Table S5; Table S6; Figure S6; Figure S7), although the difference in neonatal functional connectivity between the resilient and intermediate groups was were no longer significant (*p*=0.08). In addition, the resilient subgroup displayed larger neonatal relative volume of the right insula compared to the intermediate subgroup. For more details, please refer to the Supplemental Information.

## 4. Discussion

Using an integrative clustering approach, we identified subgroups of VPT children with distinct neurodevelopmental profiles. We described a two-cluster solution, showing a resilient subgroup with comparably favorable childhood behavioral and cognitive outcomes and increased cognitive stimulation at home, and a second, at-risk subgroup, with poor childhood behavioral and cognitive outcomes and high neonatal clinical risk. We also described a three-cluster solution, showing two subgroups largely characterized by the profiles observed in the two-cluster solution, as well as a newly emerging third intermediate subgroup with a childhood behavioral and cognitive profile intermediate between the resilient and the at-risk subgroups. Nuanced differences in socio-demographic, neonatal clinical and early brain measures appeared upon comparing subgroups from the three-cluster solution. Notably, the resilient subgroup displayed larger fronto-limbic brain regions and increased functional connectivity at term compared to the intermediate subgroup. The at-risk subgroup showed widespread white matter microstructural alterations in fronto-temporo-limbic tracts compared to the intermediate subgroup. Furthermore, the resilient subgroup had a more cognitively stimulating childhood home environment compared to the at-risk and intermediate subgroups, while the intermediate subgroup had the lowest clinical risk. Together, these findings highlight the potential value of neonatal structural and functional brain measures as useful biomarkers of later childhood outcomes in distinct VPT subgroups, as well as the importance of a supportive home environment to foster child development.

In the at-risk subgroup from the two-cluster solution, poorer childhood socio-emotional, executive function, IQ, mental health and temperament outcomes may have been driven by a combination of both higher clinical risk at birth and a less stimulating childhood home environment when compared to the resilient subgroup. Previous studies in VPT children have shown cognitively stimulating parenting to be positively associated with improved socio-emotional processing and cognitive outcomes at 2 years of age (^55^) and reduced psychopathology and executive function difficulties at 4-7 (^49^). A cognitively stimulating home environment also differentiated between psychiatric profiles at 5 (^8^). Moreover, increased neonatal clinical risk in the at-risk subgroup is consistent with previous findings, showing that perinatal medical complications following VPT birth may lead to increased behavioral and developmental problems (^15,16,56^). The resilient and at-risk subgroups, however, did not differ in any neonatal brain measures investigated, suggesting that there may be additional non-measured variables underlying different childhood outcomes, that need further investigation.

To further parse heterogeneity in VPT children, we also explored a three-cluster solution. These analyses showed that two subgroups mostly reflected the profiles seen in the two-cluster solution: 1) a resilient subgroup with high levels of childhood cognitive stimulation at home and 2) an at-risk subgroup with high levels of neonatal clinical risk. A third subgroup with intermediate childhood behavioral and cognitive profiles also emerged, in which childhood psychopathology, temperament and cognitive outcomes were poorer than those observed in the resilient subgroup, but more favorable than those observed in the at-risk subgroup. Intriguingly, the intermediate subgroup exhibited the lowest neonatal clinical risk compared to the other two subgroups, with a greater proportion of infants receiving no neonatal mechanical ventilation, CPAP or TPN and with higher median GA at birth. However, the intermediate subgroup also had higher environmental risk, namely reduced childhood cognitively stimulating home environment compared to the resilient subgroup and higher neonatal socio-demographic deprivation compared to both the at-risk and resilient subgroups. These findings suggest that developmental outcomes do not result from a single causal pathway and are best studied in a multidimensional space; for example, clinical risk, which has been linearly correlated with developmental outcomes in previous studies (^16,56^), ought to be investigated together with other factors that may influence development, i.e., environmental risk.

The at-risk compared to the intermediate subgroup showed widespread alterations in white matter microstructure (lower FA and higher MD) in the fornix, corpus callosum, corticospinal tract, inferior longitudinal, inferior fronto-occipital and uncinate fasciculi. The at-risk subgroup had also the highest neonatal clinical risk, hence the observed white matter changes are likely to be associated with preterm-related neonatal complications (^12,57,58^). White matter alterations in fronto-temporo-limbic tracts, including those observed here, have been previously associated with poorer cognitive outcomes (^59–64^). They have also been implicated in emotion processing (^65–67^) and psychiatric disorders including depression and schizophrenia (^68–70^). The intermediate subgroup, conversely, had the lowest neonatal clinical risk, and higher FA/lower MD values in fronto-temporo-limbic tracts compared to the at-risk subgroup. These findings led us to speculate that having relative fewer neonatal clinical complications, and hence fewer preterm-related white matter alterations, may contribute to these children’s more favorable cognitive, socio-emotional and behavioral outcomes compared to the at-risk subgroup. In line with this interpretation, findings in neurodegenerative conditions, suggest that ‘brain reserve’, e.g., high cortical thickness, is able to diminish the impact of brain pathology and delay the manifestation of overt clinical symptoms (^71^).

Children in the resilient subgroup exhibited higher prosocial behavior and empathy scores, as well as fewer childhood externalizing and internalizing symptoms and executive function difficulties compared to the intermediate and at-risk subgroups. They also showed lower childhood negative affectivity scores, referring to the expression of dysregulated negative emotions and increased sensitivity in response to surrounding stimuli (^72,73^). While the resilient group showed no significant brain differences compared to the at-risk subgroup, we speculate that the combination of two protective factors, an enriching home environment and lower neonatal clinical risk, may have contributed to attenuating the expression of the behavioral and cognitive difficulties associated with VPT birth. These findings also support the idea of multi-finality, whereby individuals with no overt brain differences at term may display distinct behavioral outcomes later in childhood.

Compared to the intermediate subgroup, however, the resilient subgroup displayed larger relative volumes in the left insular and bilateral orbitofrontal cortices and increased functional connectivity in an overlapping left orbitofrontal region at term, years before the behavioral and cognitive childhood outcomes were assessed. These findings could be interpreted in terms of a more advanced maturation of the fronto-limbic network in the resilient subgroup, as orbitofrontal functional connectivity and insular cortical microstructure and morphology have been positively associated with GA at birth and PMA at scan (^74,75,76^). Key nodes of the fronto-limbic network are the orbitofrontal cortex and the insula, which are structurally connected (^77^). The orbitofrontal cortex is involved in the top-down regulation of goal-oriented executive functions and socio-emotional processing (^78,79^) and the insula is important for regulating internal processes, including emotional responses to external stimuli (^80^). Structural alterations in orbitofrontal cortex and insula have been associated with emotion dysregulation (^81^) and with higher externalizing behaviors (^82^).

The orbitofrontal cortex undergoes rapid neurodevelopmental changes throughout the first year of life and its development is sensitive to environmental stimuli, such as early life stress (^83,84^). Individuals with a history of physical abuse (^85^) and VPT infants exposed to painful procedures (^86^) both show reduced orbitofrontal volumes in childhood. Furthermore, alterations in orbitofrontal connectivity and gyrification have been associated with social processing impairments in VPT children (^87^) and with executive function difficulties in extremely preterm (EPT; <28 weeks’ gestation) adolescents (^88^), respectively. Smaller insular volumes have been associated with worse emotion regulation skills (^89^) and weaker insular functional connectivity with decreased empathic responses (^90^). In the late preterm period, the insula becomes a key hub region (^91^) and a major source of transient bursting events that support brain maturation (^92^). A more mature fronto-limbic network may have therefore supported a favorable development of emotion regulation capacity, cognition, and behavior (^93,94^), resulting in the resilient subgroup exhibiting lower externalizing and internalizing symptoms, increased empathy, emotion regulation abilities and executive function skills in childhood.

This study demonstrates that it is possible to parse heterogeneity in VPT children in a meaningful way. We show that protective brain maturational patterns in the neonatal period may contribute to a more resilient behavioral profile in childhood. This is encouraging, as the preterm brain is susceptible to neuroplastic changes in response to behavioral and environmental interventions, both early in life and later in childhood (^95^). For example, neuroplastic changes have been observed following ‘supportive-touch’ (i.e., skin-to-skin contact or breastfeeding’) (^96^), maternal sensitivity training (^97^), visual stimulus cues of the mother’s face (^98^), parental praise (^99^) or music interventions in the neonatal intensive care unit (^100^). Such methods could, therefore, be used in the future to strengthen fronto-limbic circuitry to boost children’s resilience. Furthermore, our findings suggest that enriching environments may promote resilience towards more favorable behavioral outcomes. This could be done by increasing parental awareness about the importance of cognitive stimulation at home. Our findings also show that the subgroup of children with the highest neonatal clinical risk exhibit the poorest outcomes, highlighting the need to develop and implement targeted interventions for the most clinically vulnerable VPT children.

It is worth noting that the median outcome scores (IQ, BRIEF, SRS-2 and SDQ) for our three subgroups were within normative ranges and below clinical thresholds, even for the at-risk subgroup. Subthreshold psychiatric symptoms have been reported in other at-risk subgroups of VPT children (^9,8^), and have also been associated with an increased risk of developing psychiatric disorders later in life (^101^). In this context, subthreshold psychiatric symptoms may represent transdiagnostic traits that would remain undetected, and therefore untreated, if considered in a purely clinically diagnostic context, highlighting the importance of addressing psychopathology dimensionally (^102,103^).

Strengths of this study include a fairly large sample size and a rich longitudinal dataset with clinical data from birth, neonatal multi-modal MRI imaging at term and behavioral follow-up in early childhood. However, a limitation of this study is that the VPT participants included in our analyses (n=198) had a relative socio-demographic advantage than the initial baseline cohort (n=511), which may limit the generalizability of our findings to a portion of the socio-demographic spectrum. Future studies must take extra caution when interpreting such results and make increased efforts to recruit more diverse participant samples. Additional limitations to consider include the use of parental reports for most child behavioral measures, except IQ, which could lead to common method variance bias (^104^). The lack of information on familial cognitive outcomes and psychiatric history, which are heritable traits (^105^), prevents us from estimating trait heritability. Moreover, the small to moderate effect sizes reported for neonatal brain differences between subgroups may limit their immediate clinical meaningfulness or translatability into clinical practice. However, the fact that these brain differences only emerge after subdividing the sample into more refined and homogenous subgroups (C=3 vs C=2) highlights the benefit of using advanced clustering approaches such as SNF.

Sensitivity analyses including one sibling only from each twin/triplet set mostly replicated the main findings, showing similar early brain patterns as well as cognitive, neonatal clinical, social, and childhood behavioral profiles for both two-and three-cluster solutions, suggesting that the effects seen here are not biased by the presence of multiple pregnancy siblings in the main analyses.

In summary, using an integrative clustering approach, we were able to stratify VPT children into distinct multidimensional subgroups. A subgroup of VPT children at risk of experiencing behavioral and cognitive difficulties was characterized by high neonatal clinical complications and white matter microstructural alterations at term, whereas a resilient subgroup, with comparably favorable childhood behavioral outcomes, was characterized by increased childhood cognitive stimulation at home and larger and functionally more connected fronto-limbic brain regions at term. These results highlight a potential application of precision psychiatry, to enable meaningful inferences to be made at the individual level. Patterns of fronto-limbic brain maturation may be used as image-based biomarkers of outcomes in VPT children, while promoting enriching environments may foster more optimal behavioral outcomes. Risk stratification following VPT birth could, therefore, guide personalized behavioral interventions aimed at supporting healthy development in vulnerable children.

## Acknowledgements

We would like to thank participant families and children that took part in this study as well as research radiographers, administrative staff and clinical personnel that made this work possible.

This work was supported by the Medical Research Council (MRC), UK [grant numbers: MR/K006355/1, MR/S026460/1 and MR/N013700/1], Action Medical Research and Dangoor Education [grant number: GN2606], King’s College London member of the MRC Doctoral Training Partnership in Biomedical Sciences and Biotechnology and Biological Sciences Research Council [grant number: BB/J014567/]. This study uses data acquired during independent research funded by the National Institute for Health Research (NIHR) Programme Grants for Applied Research Programme [grant number: RP-PG-0707-10154] and the research is supported by the NIHR Biomedical Research Centre, Guy’s and St Thomas’ NHS Foundation Trust and King’s College London, the NIHR Clinical Research Facility, Guy’s and St Thomas’ and the MRC Centre for Neurodevelopmental Disorders.

The authors acknowledge use of the research computing facility at King’s College London, *Rosalind* (https://rosalind.kcl.ac.uk), which is delivered in partnership with the National Institute for Health Research (NIHR) Biomedical Research Centres at South London & Maudsley and Guy’s & St. Thomas’ NHS Foundation Trusts, and part-funded by capital equipment grants from the Maudsley Charity [award 980] and Guy’s & St. Thomas’ Charity [TR130505]. The views expressed are those of the author(s) and not necessarily those of the NHS, the NIHR, King’s College London, or the Department of Health and Social Care.

## Data availability

Access to the dataset supporting this article can be made available upon request from the corresponding author.

*For the purpose of open access, the author has applied a Creative Commons Attribution (CC BY) licence to any Author Accepted Manuscript version arising*.

