## Supplemental Information document for "Parsing brain-behavior heterogeneity in very preterm born children using integrated similarity networks"

**Running title:** Brain-behavior stratification of preterm children

Laila Hadaya (MSc)^1,2^, Konstantina Dimitrakopoulou (PhD)^3^, Lucy Vanes (PhD)^4^, Dana Kanel (PhD)^1,2^, Sunniva Fenn-Moltu (MSc)^1,5^, Oliver Gale-Grant (MD)^1,5,6^, Serena J Counsell (PhD)^1^, A David Edwards (FMedSci)^1^, Mansoor Saqi (PhD)^3^, Dafnis Batalle (PhD)^1,5^, Chiara Nosarti (PhD)^1,2^

^1^Centre for the Developing Brain, Department of Perinatal Imaging and Health, King’s College London, London, United Kingdom. ^2^Department of Child and Adolescent Psychiatry, Institute of Psychiatry Psychology and Neuroscience, King’s College London, London, United Kingdom. ^3^Translational Bioinformatics Platform, NIHR Biomedical Research Centre, Guy's and St. Thomas' NHS Foundation Trust and King's College London, London, United Kingdom. ^4^Centre for Neuroimaging Sciences, Institute of Psychiatry Psychology and Neuroscience, King’s College London, London, United Kingdom. ^5^Department of Forensic and Neurodevelopmental Sciences, Institute of Psychiatry Psychology and Neuroscience, King’s College London, London, United Kingdom. ^6^MRC Centre for Neurodevelopmental Disorders, King’s College London, United Kingdom.

**Correspondence to**: Chiara Nosarti, Centre for the Developing Brain, Department of Perinatal Imaging and Health, School of Biomedical Engineering & Imaging Sciences, King’s College London, First Floor South Wing, St Thomas’ Hospital, London SE1 7EH

**MRI acquisition parameters**

Images were acquired with the following parameters: T2-weighted fast-spin echo sequences: TR = 8670 ms, TE = 160 ms, flip angle = 90°, slice thickness = 1 mm, FOV = 220 x 220 mm, matrix = 256 x 256, voxel size = 0.86x0.86x1 mm. Single-shot echo-planar diffusion MRI was acquired in the transverse plane in 32 non-collinear directions: (TR = 8000 ms; TE = 49 ms, slice thickness = 2 mm, FOV = 224x224 mm, matrix = 128x128, voxel size = 1.75x1.75x2 mm, b-value = 750 s/mm^2^, SENSE factor 2). T2-gradient echo-planar imaging at rest: TR= 1.5 s, ET = 45 ms, flip angle = 90°, 256 volumes, slice thickness= 3.25 mm, in-plane resolution= 2.5x2.5 mm, 22 slices, scan duration= 6.4 mins. Infants received MRI while asleep and 87% were sedated (prior to MRI) with 25-50 mg/kg chloral hydrate. Neonatal earmuffs (MiniMuffs, Natus Medical Inc., San Carlos, CA, USA) and earplugs molded from silicone-based putty (President Putty, Coltene Whaledent, Mahwah, NJ, USA) were used for ear protection.

**Figure S1. Study sample flowchart**

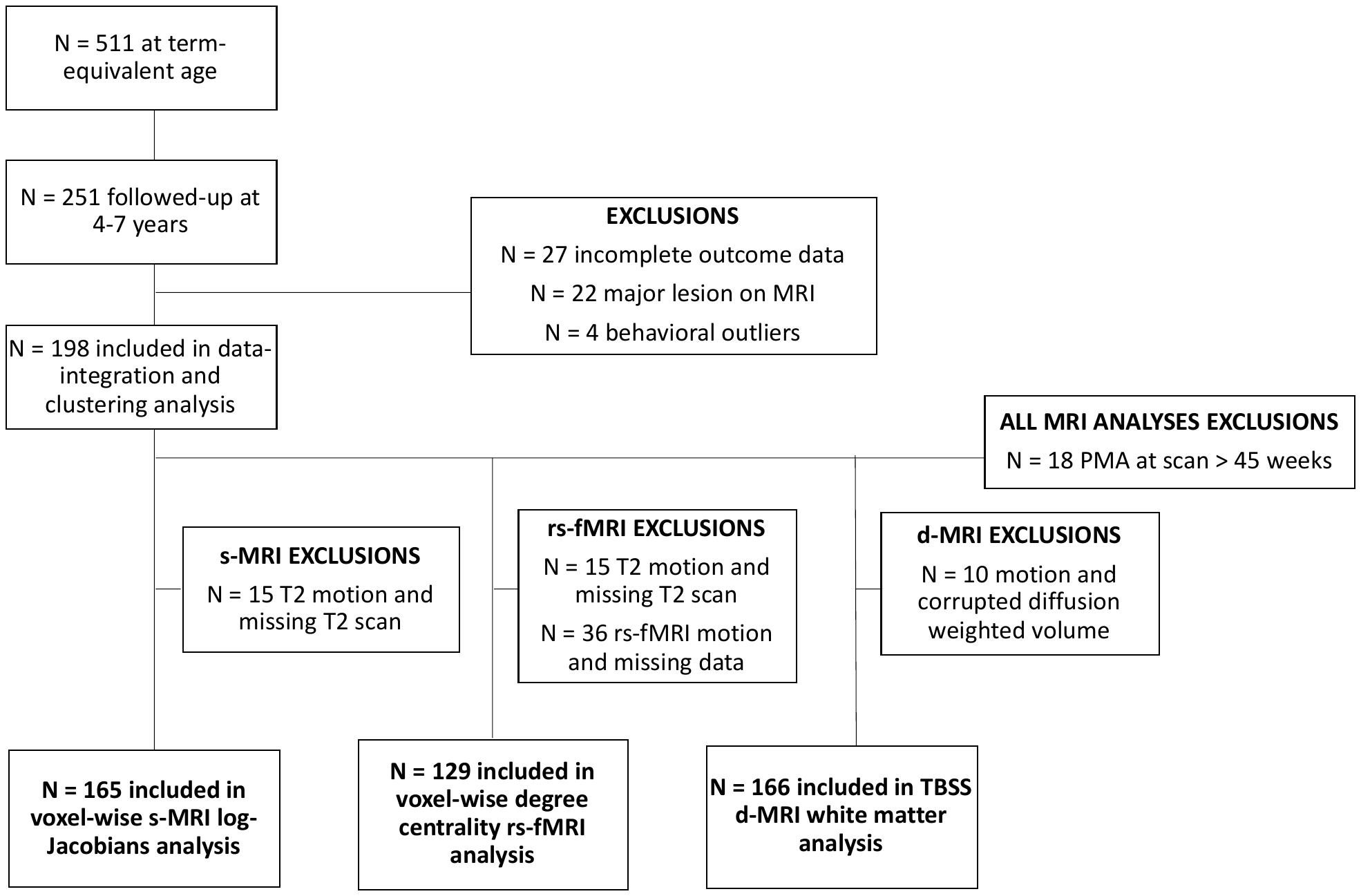

**Ethnicity in Table 1 was grouped according to the Office of National Statistics classifications (2016).**

The groups were characterised accordingly: White (English/Welsh/Scottish/Northern Irish/British, Irish, Any other White background); Mixed/Multiple ethnic groups (White and Black Caribbean, White and Black African, White and Asian, Any other Mixed/Multiple ethnic background); Asian/Asian British (Indian, Pakistani, Bangladeshi, Chinese, Any other Asian background); Black/African/Caribbean/Black British (African, Caribbean, Any other Black/African/Caribbean background); Other ethnic group (Arab, Any other ethnic group).

**Data integration and clustering pipeline**

An adaptation of the *ExecuteSNF.CC* function (^1^) was used to generate distance and similarity matrices and run both integration and consensus clustering algorithms on a combination of mixed (type 1) and numeric (type 2 and 3) data types. Code can be accessed here: <https://github.com/lailahadaya/preterm-ExecuteSNF.CC>

***Data integration***

Similarity Network Fusion (SNF) is a message passing theory method, which updates the final fused matrix over a series of iterations (specified by hyperparameter T; default T=20), increasing the signal-to-noise ratio by updating the final fused matrix with each iteration and discarding weak and inconsistent connections (edges) between subjects (nodes) across data types and maintaining stronger or consistent edges across data types with a neighborhood size of K (^2,3^)

***Clustering***

The “consensus clustering” resultant subgroups obtained in step 2 (called *consensus clusters*) were drawn from an agglomerative hierarchical clustering approach, which partitioned individuals into subgroups based on the pairwise *consensus value* (i.e., the proportion of times each two individuals co-clustered together across the several clustering attempts) (^4,5^). Silhouette width scores were used to measure clustering quality for each subject/node within the network, with values ranging from -1 to 1, where larger values indicate a subject is closer to other subjects within the same cluster than to subjects in other clusters and values closer to -1 indicate misclassification.

**Estimation of number of clusters**

To estimate the optimal number of clusters (between C=2 and C=10), Eigengap and Rotation Cost heuristics were run for each combination of K-alpha hyperparameters. A two-cluster solution (C=2) was estimated to be the optimal number of clusters 60/60 times (30/30 times as the *best* solution using Eigengap and 30/30 times as the *best* solution using Rotation Cost), followed by C=3 (25/60 times; 13/30 times as the *second best* solution using Eigengap and 12/30 times as the *second best* using Rotation Cost) and C=4 (31/60 times; 16/30 times as the *second best* solution using Eigengap and 15/30 times as the *second best* solution using Rotation Cost) (Figure S2A).

The clustering steps were repeated for C=2, C=3, C=4, consensus matrices were constructed (Figure S2B; S2C) and Silhouette width scores were calculated using the consensus matrices (Figure S2D). The sample was clustered into C=2 and C=3 subgroups and analyzed for phenotypic differences, as the average Silhouette width scores were highest and had the least number of negative Silhouette width values (Figure S2D) and as consensus matrices also reflected higher proportions of co-clustering over multiple runs for both C=2 and C=3 compared to C=4.

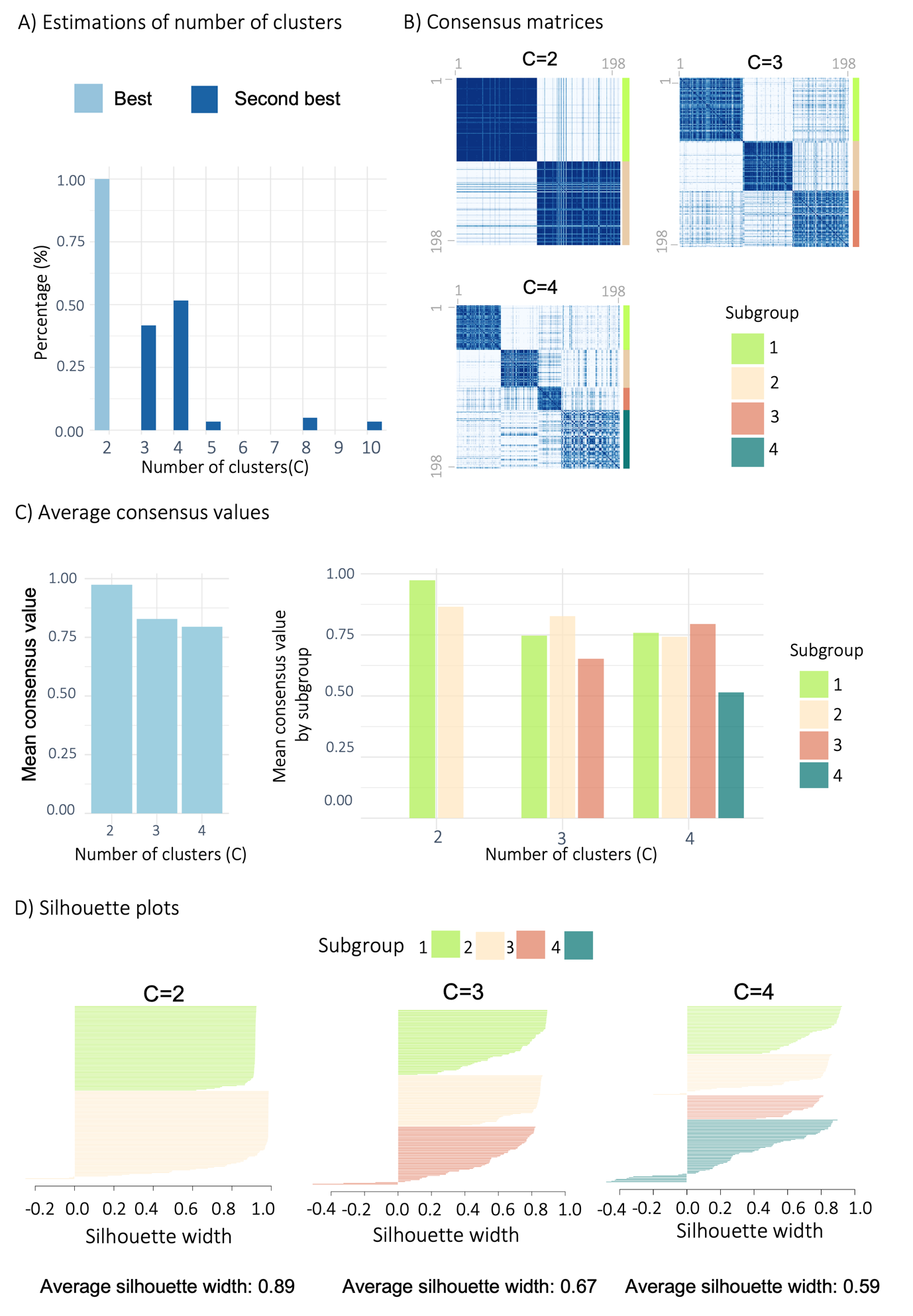

**Figure S2. Estimating the optimal number of clusters**. **A)** the number of times each number was selected as the most optimal number of clusters (best; pale blue), and second most optimal number of clusters (second best; blue) for the 30 combinations of K-alpha hyperparameters using Eigengap and Rotation Cost. **B)** Consensus matrices were calculated after clustering into C=2, C=3 and C=4 respectively. Each consensus matrix (the sum of times each pair of subjects co-clustered divided by the sum of times each pair of subjects were both present in a given iteration’s subsample) reflects the proportion of times each pair of subjects co-clustered. Darker blue represents higher co-clustering proportions and lighter colors reflect lower proportions. Final clustering results are shown for the four subgroups (green = subgroup 1; beige = subgroup 2; pale pink = subgroup 3; teal = subgroup 4). **C)** Left: mean consensus pairwise values (mean proportion of times each pair of subjects co-clustered) for each number of clusters (C=2, C=3 and C=4), Right: mean pairwise consensus values for pairs of subjects co-clustering in each subgroup for C=2, C=3 and C=4. **D)** Silhouette width values for each subgroup after clustering into C=2, C=3, C=4 (left to right respectively).

As seen in Figure S3, the majority of participants in subgroup 1 from the two-cluster solution (bottom left; C=2) were preserved into subgroup 1 from the three-cluster solution (bottom right; C=3). Similarily, a large proportion of participants from subgroup 2 in C=2 (top left) were preserved into subgroup 2 from C=3 (top right). The third subgroup (subgroup 3) from C=3 consists of participants that moved from both C=2 subgroup 1 and subgroup 2. Participants in the two subgroups from the two-cluster solution (C=2 subgroup 1 and subgroup 2) that moved into a cluster with a corresponding behavioral profile in the three-cluster solution (C=3 subgroup 1 and subgroup 2) are represented by grey lines, while red lines seen represent participants that moved into the newly emerging subgroup 3.

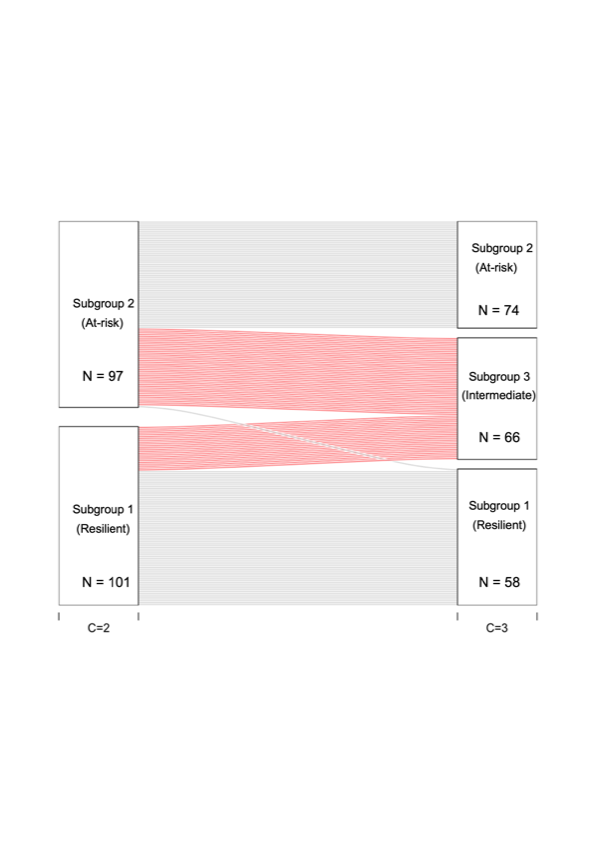

**Figure S3. Alluvial plot showing the transition of subject assignment from C=2 subgroups to C=3 subgroups.**

**Selection of in-model and out-of-model variables**

Given the prevalence of socio-emotional and executive function difficulties in VPT children, as well as the importance of both clinical and environmental risk factors in shaping these outcomes, relevant variables from these domains were included in the data integration and clustering model, in order to parse heterogeneity in the sample. EmQue empathy subscales and SRS-2 scores were used in-model as measures of socio-emotional processing, which is critical for social communication and interaction (^6–8^). The BRIEF subscales (inhibit, shift, emotional control, working memory and plan/organize) were used as measures of executive function, behavioral self-regulation and cognitive control (^9^). Four clinical variables (days TPN, days CPAP, days ventilation as well as GA at birth) were included in the model based on our previous factor analysis in an overlapping cohort sample, to obtain a clinical summary score derived from 28 clinical variables, as described in (^10^). A measure of neighborhood deprivation (Index of Multiple Deprivation) was also used in-model as a measure of socio-demographic risk. Out-of-model cognitive, behavioral and environmental variables were selected in order to provide external validation of the resultant subgroup profiles. These variables were IQ, CSPS total score, temperamental traits (CBQ negative affectivity, surgency and effortful control scores), which reflect the ability to regulate emotions and behaviors in responses to emotional stimuli (^11^) and SDQ internalizing and externalizing symptom scores.

A correlation plot summarising Spearman Rho correlation coefficients between in-model and out-of-model variables is shown below (Figure S4). The correlations between the out-of-model variables with the in-model variables range from weak to moderate.

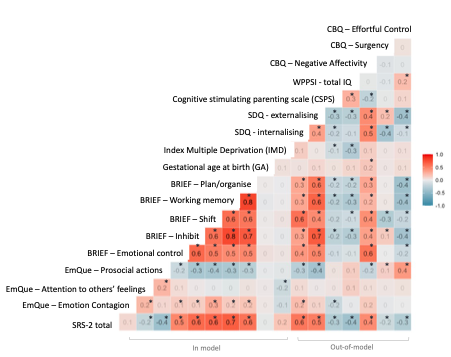

**Figure S4 Correlation plot of in-model and out-of-model variables.** Numbers denote Spearman R values for each pair of variables, which are depicted with a color gradient (see colorbar). P-values are denoted by asterisks whereby *=*p*<0.05 (uncorrected).

**Post-hoc analysis – clustering based on neonatal socio-demographic and clinical risk factors only**

In order to further demonstrate the benefit of using a cluster solution generated by an integrative clustering approach in comparison to that from clustering only one data type, we ran a post-hoc analysis where we clustered the cohort based on only their neonatal socio-demographic and clinical risk factors (i.e., single data type). As expected, results changed substantially (see Figure S5). The alluvial plots indicate the allocation of each subject (where each subject is represented by a red line) within subgroups created after clustering with all data types (left) or with only the socio-demographic and clinical risk data type (right). We also investigated the resultant subgroups for differences in childhood outcomes and found no significant differences for C=2 and C=3 (all *p*s>0.05). These results further highlight the importance of integrating heterogenous data types which capture information from different domains.

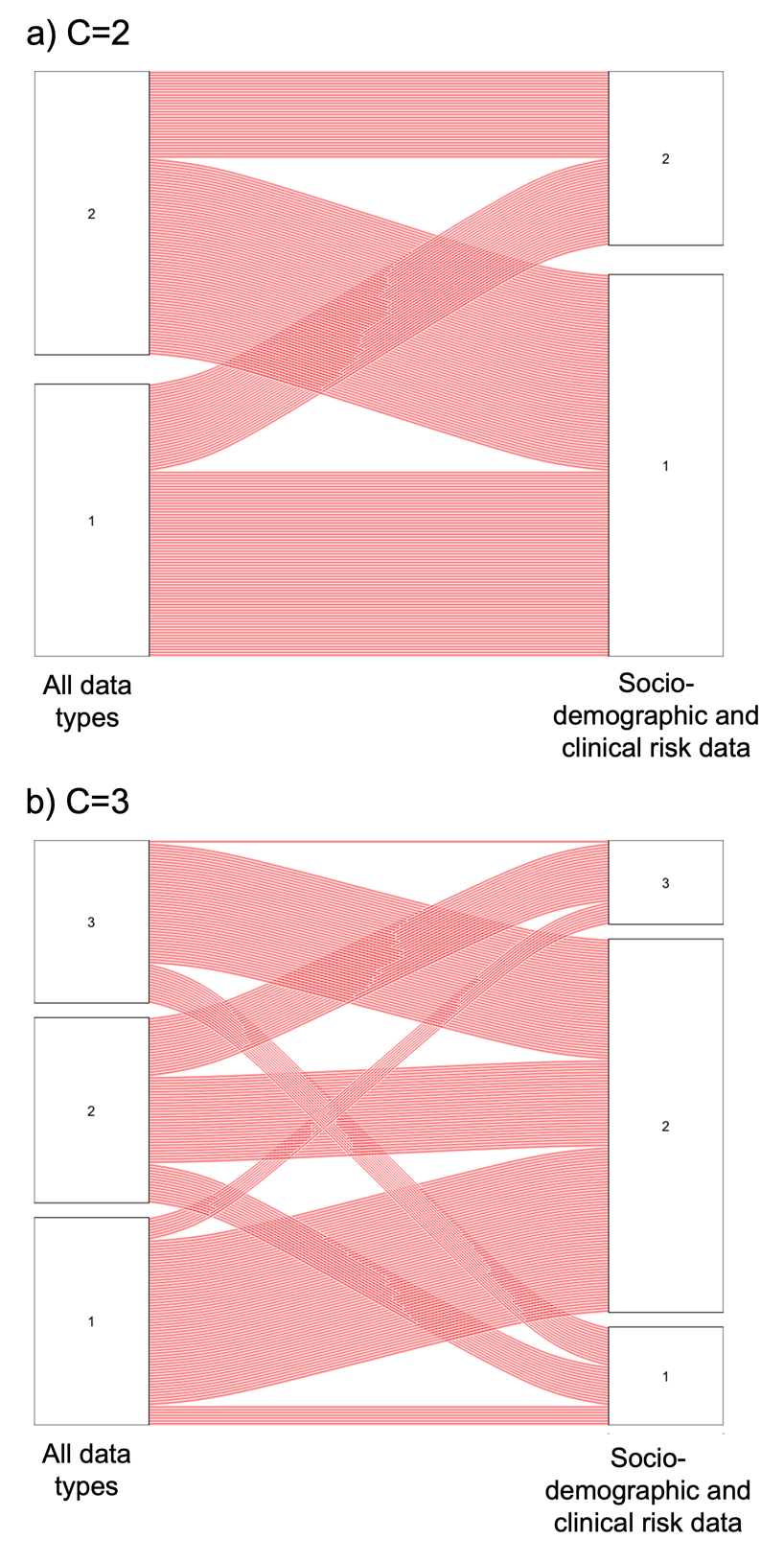

**Figure S5. Alluvial plot indicating allocation of participants in subgroups after implementing integrative clustering based on all data types (left) and clustering of only the socio-demographic and clinical risk data type (right) for a) two clusters (C=2) and b) three clusters (C=3).**

**MRI pre-processing and analyses**

***Diffusion MRI image pre-processing and Tract Based Spatial Statistics***

Diffusion MRI (d-MRI) pre-processing steps are described in our previous work (^10^,^12^). To briefly summarize, images were visually checked for motion artefacts and corrupted volumes were excluded. All d-MRI datasets that were included in the analysis had a total of 5 volumes or fewer excluded. BET (version 2.1; http://fsl.fmrib.ox.ac.uk/fsl/fslwiki/BET) (^13^,^14^) was used to extract non-brain tissue from the data and *eddy_current* to correct for eddy current artefacts (^15^). FSL’s *dtifit* was used to fit the tensor model (FMRIB, Oxford, http://fsl.fmrib.ox.ac.uk).

Image registration was performed using DTI-TK and integrated within the TBSS pipeline to produce a population specific DTI template. From this template a mean FA map was derived and then thinned by perpendicular non-maximum suppression to create a mean FA skeleton. A FA threshold of ≥ 0.15 was used to limit the inclusion of voxels with high inter-subject variability and non-white matter voxels. FA and MD maps were projected onto this skeleton prior to voxel-wise statistical analysis.

***Tensor Based Morphometry processing***

Two input modalities (T2-weighted images and T2-weighted image tissue type segmentations) were registered to a study-specific T2-weighted template using the ANTS software multimodal Symmetric Normalisation (SyN) (^16^) algorithm as described in Lautarescu et al. (^17^).

Deformation tensor field gradients (i.e., log Jacobian determinant maps) were then computed from the resultant T2-weighted deformation tensor fields (i.e., warps) of the non-linear transformations. Jacobian determinant map values reflect the degree of contraction or expansion a voxel undergoes following the non-linear transformation from native to template space (^18^). The logarithm Jacobian maps were smoothed (4 mm FWHM Gaussian filter) to improve the signal to noise ratio. Data were resampled from 0.5 to 1 mm^3^ isotropic voxel size to decrease computation and memory load.

***Functional MRI image pre-processing and motion-correction***

Functional images were pre-processed as in Ball et al. (^19^). In summary, images with visible motion artefacts were excluded after visual inspection. Single-subject independent component analysis (ICA) with automatic dimensionality estimation was applied to each individual’s dataset using FSL MELODIC (^20^), following removal of the first 6 volumes (to allow for T1 equilibration), motion correction using MCFLIRT, and high-pass filtering (125 s cutoff, 0.008 Hz). Following ICA, FSL FIX (^21^) was applied for automatic denoising and artefact removal. A population-specific neonatal template with tissue priors was used for standard-space masking (^22^), and the FIX algorithm was trained on hand-classified fMRI datasets from 40 preterm infants aged 28-44 weeks, collected on the same scanner (including both low-motion and high-motion subjects; for more details see supplemental materials of (^19^)). After components were classified as either signal or noise, the unique variance of each noise component, as well as the full variance of the motion parameters and derivatives, were regressed out of the data (^23,24^). Standardized DVARS, a framewise data quality index (^25^), was calculated before and after applying FIX and significantly improved after FIX clean-up (t(315)=9.01, *p*<0.001). Finally, FSL Motion Outliers was applied to each dataset to identify remaining volumes that were corrupted by large motion, and subjects with more than two standard deviations above the mean number of corrupted volumes were removed. This resulted in a final fMRI sample of 298 infants. Of these, 129 with complete follow-up data were included in further analysis.

Cleaned functional images from this sample were resampled to 2 mm^3^ isotropic voxels, registered to the study-specific T2-weighted template using boundary-based registration, and smoothed with a 4 mm full-width half-maximum Gaussian kernel.

**Table S1. Subgroup sample sizes for each imaging modality**

| **Cluster solution** | **Subgroup** | **d-MRI sample** | **s-MRI sample** | **rs-fMRI sample** | **Total sample** |
| --- | --- | --- | --- | --- | --- |
| **C=2** | Resilient | 80 | 82 | 63 | 97 |
|  | At-risk | 86 | 83 | 66 | 101 |
| **C=3** | Resilient | 60 | 61 | 48 | 74 |
|  | At-risk | 52 | 48 | 38 | 58 |
|  | Intermediate | 54 | 56 | 43 | 66 |
| **Total sample** | | 166 | 165 | 129 | 198 |

**Table S2. Two-cluster solution profiles using in-model and out-of-model variables**

|  | Variables | Subgroup 1 | Subgroup 2 | p-value | Effect size |
| --- | --- | --- | --- | --- | --- |
| In-model Variables | **SRS-2** | 22.00 (5.00) | 43.00 (5.75) | <0.001 | -0.79 |
|  | **EmQue: Emotion Contagion** | 0.17 (0.33) | 0.33 (0.67) | 0.001 | -0.26 |
|  | **EmQue: Attention to others’ emotions** | 1.43 (0.57) | 1.29 (0.43) | 0.933 | 0.01 |
|  | **EmQue: Prosocial actions** | 1.33 (0.50) | 0.83 (0.50) | <0.001 | 0.53 |
|  | **BRIEF: Emotion control** | 13 (4.00) | 16.50 (4.43) | <0.001 | -0.52 |
|  | **BRIEF: Inhibit** | 22.00 (5.00) | 28.50 (8.75) | <0.001 | -0.73 |
|  | **BRIEF: Shift** | 11.00 (2.00) | 15.00 (5.00) | <0.001 | -0.81 |
|  | **BRIEF: Working memory** | 21.00 (5.00) | 29.00 (8.75) | <0.001 | -0.81 |
|  | **BRIEF: Plan & organize** | 14.00 (3.00) | 17.50 (3.00) | <0.001 | -0.73 |
|  | **GA** | 30.29 (4.00) | 29.93 (4.43) | 0.590 | 0.04 |
|  | **IMD at birth** | 15.99 (20.68) | 15.15 (14.36) | 0.888 | 0.01 |
|  | **Days TPN** (0:1:2), n= | 33:53:11 | 35:45:21 | 0.153  (X^2^=3.76) | V=0.14 |
|  | **Days ventilation** (0:1:2), n= | 58:35:4 | 43:39:10 | 0.002 | V=0.25 |
|  | **Days CPAP** (0:1:2), n= | 23:55:19 | 10:70:21 | 0.031  (X^2^=6.94) | V=0.18 |
| Out-of-model variables | **SDQ total** | 6.00 (5.00) | 11.00 (5.75) | <0.001 | -0.73 |
|  | **SDQ: Internalizing** | 1.00 (3.00) | 4.00 (3.75) | <0.001 | -0.56 |
|  | **SDQ: Externalizing** | 4.00 (3.00) | 8.00 (5.00) | <0.001 | -0.61 |
|  | ^$^**CSPS** | 18.00 (3.00) | 17.00 (3.53) | 0.003 | 0.24 |
|  | ^$$^**WPPSI: Full scale IQ** | 110.00 (17.50) | 104.50 (26.00) | 0.026 | 0.18 |
|  | **CBQ: Negative affectivity** | 3.67 (1.10) | 4.42 (00.96) | <0.001 | -0.49 |
|  | **CBQ: Surgency** | 5.00 (0.92) | 4.67 (1.31) | 0.057 | 0.16 |
|  | **CBQ: Effortful control** | 5.50 (0.92) | 5.17 (1.23) | 0.002 | 0.25 |
|  | **Corrected age at assessment: years** | 4.67 (0.82) | 4.59 (0.58) | 0.074 | 0.15 |
|  | **IMD at assessment** | 13.75 (19.58) | 15. 88 (16.51) | 0.499 | -0.057 |
|  | **Sex** (M:F), n= | 50:47 | 50:51 | 0.885 (X^2^=0.02) | V=0.02 |
|  | **Total,** n (%) | 97 (48.99%) | 101 (51.01%) | / |  |
| *Note: Median (IQR) is provided unless otherwise stated. P-values refer to results from Mann-Whitney non-parametric test. Effect sizes reported are Glass Rank Biserial Correlation unless otherwise stated, where Cramer’s V is reported for categorical variables. Days TPN (0:1:2), days ventilation (0:1:2) and days CPAP (0:1:2), correspond to the ratio of the three clinical risk categories: 0, 1 and 2. Respectively, they correspond to zero days, more than zero days but less than the top quintile, and within the top quintile. ^$^=one missing participant; ^$$^=two missing participants. Chi-squared test was used for categorical comparisons.*  *Abbreviations: BRIEF = Behavior Rating Inventory of Executive Function; CBQ = Childhood Behavioral Questionnaire; CSPS = Cognitively Stimulating Parenting Scale; CPAP = continuous positive airway pressure. EmQue = Empathy Questionnaire; GA = gestational age; IMD = Index of Multiple Deprivation; IQR = interquartile range; PMA = post menstrual age at scan; SDQ = Strengths and Difficulties Questionnaire; SRS-2 = Social Responsiveness Scale – Second Edition; TPN = total parenteral nutrition.* | | | | | |

**Table S3. Three-cluster solution profiles using in-model and out-of-model variables**

|  | Variables | Subgroup 1 | Subgroup 2 | Subgroup 3 | p-value | Effect size |
| --- | --- | --- | --- | --- | --- | --- |
| In-model Variables | **SRS-2** | 18.00 (14) | 56.50 (24.50) | 12.50 (36.00) | <0.001 | 0.55 |
|  | **EmQue: Emotion Contagion** | 0.17 (0.33) | 0.50 (0.67) | 0.17 (0.33) | 0.003 | 0.08 |
|  | **EmQue: Attention to others’ emotions** | 1.43 (0.57) | 1.29 (0.43) | 1.29 (0.57) | 0.186 | 0.01 |
|  | **EmQue: Prosocial actions** | 1.33 (0.46) | 0.83 (0.50) | 1.00 (0.33) | <0.001 | 0.24 |
|  | **BRIEF: Emotion control** | 13.00 (5.00) | 18.00 (5.00) | 14.00 (4.75) | <0.001 | 0.34 |
|  | **BRIEF: Inhibit** | 21.00 (5.00) | 31.50 (8.75) | 25.00 (6.00) | <0.001 | 0.44 |
|  | **BRIEF: Shift** | 11.00 (2.00) | 17.00 (4.75) | 13.00 (2.00) | <0.001 | 0.59 |
|  | **BRIEF: Working memory** | 21.00 (5.00) | 31.00 (8.75) | 26.00 (5.00) | <0.001 | 0.48 |
|  | **BRIEF: Plan & organize** | 13.00 (4.00) | 18.00 (4.50) | 16.00 (3.00) | <0.001 | 0.42 |
|  | **GA** | 29.36 (3.64) | 30.00 (4.79) | 30.64 (3.39) | 0.003 | 0.04 |
|  | **IMD at birth** | 13.23 (14.77) | 14.12 (9.86) | 22.08 (17.11) | 0.018 | 0.06 |
|  | **Days TPN** (0:1:2), n= | 16:48:10 | 17:28:13 | 35:22:9 | <0.001 (X^2^=19.64) | V=0.22 |
|  | **Days ventilation** (0:1:2), n= | 40:30:4 | 23:21:14 | 38:23:5 | 0.014 | V=0.19 |
|  | **Days CPAP** (0:1:2), n= | 9:45:20 | 2:40:16 | 22:40:4 | <0.001 | V=0.27 |
| Out-of-model variables | **SDQ total** | 5.00 (4.00) | 13.00 (5.75) | 8.00 (5.00) | <0.001 | 0.40 |
|  | **SDQ: Internalizing** | 1.00 (2.00) | 4.00 (4.75) | 2.00 (2.83) | <0.001 | 0.28 |
|  | **SDQ: Externalizing** | 4.00 (3.00) | 9.00 (5.00) | 6.00 (4.00) | <0.001 | 0.28 |
|  | **CSPS** | 19.00 (2.00) | 17.00 (3.00) | 18.00 (4.00) | 0.006 | 0.28 |
|  | **WPPSI: Full scale IQ** | 112.00 (16.00) | 103.50 (30.25) | 105.00 (22.50) | 0.007 | 0.07 |
|  | **CBQ: Negative affectivity** | 3.58 (1.15) | 4.50 (1.04) | 4.25 (0.98) | <0.001 | 0.20 |
|  | **CBQ: Surgency** | 4.96 (0.83) | 4.61 (1.21) | 4.75 (1.25) | 0.071 | 0.03 |
|  | **CBQ: Effortful control** | 5.67 (0.83) | 5.04 (1.33) | 5.21 (0.99) | 0.002 | 0.10 |
|  | **Corrected age at assessment: years** | 4.63 (0.80) | 4.60 (0.56) | 4.69 (1.07) | 0.78 | 0.01 |
|  | **IMD at assessment** | 12.02 (14.44) | 18.73 (17.18) | 15.15 (17.99) | 0.133 | 0.034 |
|  | **Sex** (M:F), n= | 36:38 | 29:29 | 35:31 | 0.871 (X^2^=0.28) | V=0.04 |
|  | **Total**, n (%) | 74 (37.37%) | 58 (29.29%) | 66 (33.33%) | / |  |
| *Note*: *Median (IQR) is provided unless otherwise stated. P-values refer to results from Kruskal-Wallis non-parametric* test*. Effect sizes reported are Epsilon Squared unless otherwise stated where Cramer’s V is reported for categorical variables.*  Days TPN (0:1:2), days ventilation (0:1:2) and days CPAP (0:1:2), correspond to the ratio of the three clinical risk categories: 0,1 and 2. Respectively, they correspond to zero days, more than zero days but less than the top quintile and within the top quintile. ^$^=one missing participants; ^$$^=two missing participants. Chi-squared test was used for categorical comparisons when subject count per cell was >5 and Fisher’s Exact when cell count was 5 or less.  Abbreviations: *BRIEF = Behavior Rating Inventory of Executive Function; CBQ = Childhood Behavioral Questionnaire; CSPS = Cognitively Stimulating Parenting Scale; CPAP = continuous positive airway pressure. EmQue = Empathy Questionnaire; GA = gestational age; IMD = Index of Multiple Deprivation; IQR = interquartile range; PMA = post menstrual age at scan; SDQ = Strengths and Difficulties Questionnaire; SRS-2 = Social Responsiveness Scale – Second Edition; TPN = total parenteral nutrition.* | | | | | |  |

**Table S4. Effect sizes, number of significant voxels and p-values for brain regions showing significant differences between subgroups**

| Brain measure | Contrast | Region | Number of voxels | p-value | Cohen's F effect size |
| --- | --- | --- | --- | --- | --- |
| Log-Jacobian determinant | **Resilient > intermediate** | Left insular | 553 | 0.01 | 0.52 |
|  | **Resilient > intermediate** | Left orbitofrontal | 1652 | 0.01 | 0.46 |
|  | **Resilient > intermediate** | Right orbitofrontal regions | 897 | 0.01-0.05 | 0.48 |
| Degree centrality | **Resilient > intermediate** | Left orbitofrontal | 11 | 0.04 | 0.45 |
| TBSS – FA | **Intermediate > at-risk** | Fornix, CC, CST, ILF, IFO and UF | 1370 | 0.01 | 0.08 |
| TBSS – MD | **Intermediate < at-risk** | Fornix and CC body | 93 | 0.04 | 0.05 |
| Note.  Abbreviations: CC= corpus callosum, CST= corticospinal tract, FA = fractional anisotropy, IFO = inferior fronto-occipital fasciculus, ILF=inferior longitudinal fasciculus, MD = mean diffusivity, TBSS = tract based spatial statistics, UF = uncinate fasciculus. | | | | | |

**Sensitivity analyses**

In order to account for multiple pregnancy confounding, we conducted sensitivity analyses including only one child, at random, from each set of multiple pregnancy siblings. Results for the in-model and out-of-model cognitive, behavioral, clinical and socio-demographic risk variables remained similar to results including children born from multiple pregnancy for both C=2 (Table S5) and C=3 (Table S6). Sensitivity analysis results showed that the log-Jacobian brain differences observed between the subgroups were largely preserved (Figure S6), with the resilient subgroup having larger brain volumes in the left insula and bilateral orbitofrontal cortices compared to the intermediate group (Figure S6A) and the intermediate group showing higher FA and lower MD in white matter tracts compared to the at-risk subgroup (Figure S6C). Results of sensitivity analysis also showed an additional result that was not observed in the full sample: the resilient subgroup displayed larger brain volumes in the right insula compared to the intermediate subgroup. However, the functional connectivity degree centrality was no longer significant, with *p*=0.08 (Figure S6B). We believe this may have been due to a loss in power, as a result of the reduction in sample size.

**Table S5. Results of sensitivity analysis, including only one child from each set of multiple pregnancy siblings, of the two-cluster solution profiles using in-model and out-of-model variables**

|  | Variables | Subgroup 1 | Subgroup 2 | p-value |
| --- | --- | --- | --- | --- |
| In-model Variables | **SRS-2** | 22.00 (5) | 43.00 (5.75) | <0.001 |
|  | **EmQue: Emotion Contagion** | 0.17 (0.33) | 0.33 (0.67) | 0.005 |
|  | **EmQue: Attention to others’ emotions** | 1.43 (0.57) | 1.29 (0.43) | 0.658 |
|  | **EmQue: Prosocial actions** | 1.33 (0.50) | 0.83 (0.50) | <0.001 |
|  | **BRIEF: Emotion control** | 13 (4.00) | 16.50 (4.43) | <0.001 |
|  | **BRIEF: Inhibit** | 22.00 (5.00) | 28.50 (8.75) | <0.001 |
|  | **BRIEF: Shift** | 11.00 (2.00) | 15.00 (5.00) | <0.001 |
|  | **BRIEF: Working memory** | 21.00 (5.00) | 29.00 (8.75) | <0.001 |
|  | **BRIEF: Plan & organize** | 14.00 (3.00) | 17.50 (3.00) | <0.001 |
|  | **GA** | 30.29 (4.00) | 29.93 (4.43) | 0.315 |
|  | **IMD** | 15.99 (20.68) | 15.15 (14.36) | 0.968 |
|  | **Days TPN** (0:1:2), n= | 27:43:11 | 31:38:21 | 0.197 (X^2^=3.25) |
|  | **Days ventilation** (0:1:2), n= | 49:28:4 | 36:36:18 | 0.004 |
|  | **Days CPAP** (0:1:2), n= | 18:48:15 | 9:61:20 | 0.091 (X^2^=4.80) |
| Out-of-model variables | **SDQ total** | 6.00 (5.00) | 11.00 (5.75) | <0.001 |
|  | **SDQ: Internalising** | 1.00 (3.00) | 4.00 (3.75) | <0.001 |
|  | **SDQ: Externalising** | 4.00 (3.00) | 8.00 (5.00) | <0.001 |
|  | ^$^**CSPS** | 18.00 (3.00) | 17.00 (3.53) | 0.006 |
|  | ^$$^**WPPSI: Full scale IQ** | 110.00 (17.50) | 104.50 (26.00) | 0.019 |
|  | **CBQ: Negative affectivity** | 3.67 (1.10) | 4.42 (00.96) | 0.058 |
|  | **CBQ: Surgency** | 5.00 (0.92) | 4.67 (1.31) | <0.001 |
|  | **CBQ: Effortful control** | 5.50 (0.92) | 5.17 (1.23) | 0.012 |
|  | **Corrected age at assessment: years** | 4.67 (0.82) | 4.59 (0.58) | 0.082 |
|  | **IMD at assessment** | 13.75 (19.65) | 15.75 (16.76) | 0.553 |
|  | **Sex** (M:F), n= | 43:38 | 43:47 | 0.589 (X^2^=0.29) |
|  | **Total**, n (%) | 81 (47.37%) | 90 (52.63%) | / |
| *Note: Median (IQR) is provided unless otherwise stated. P-values refer to results from Mann-Whitney non-parametric test. Days TPN (0:1:2), days ventilation (0:1:2) and days CPAP (0:1:2), correspond to the ratio of the three clinical risk categories: 0,1 and 2. Respectively, they correspond to zero days, more than zero days but less than the top quintile and within the top quintile. ^$^=one missing participant; ^$$^=two missing participants. Chi-squared test was used for categorical comparisons when subject count per cell was >5 and Fisher’s Exact when cell count was 5 or less.*  *Abbreviations: BRIEF = Behavior Rating Inventory of Executive Function; CBQ = Childhood Behavioral Questionnaire; CSPS = Cognitively Stimulating Parenting Scale; CPAP = continuous positive airway pressure. EmQue = Empathy Questionnaire; GA = gestational age; IMD = Index of Multiple Deprivation; IQR = interquartile range; PMA = post menstrual age at scan; SDQ = Strengths and Difficulties Questionnaire; SRS-2 = Social Responsiveness Scale – Second Edition; TPN = total parenteral nutrition.* | | | | |

**Table S6. Results of sensitivity analysis, including only one child from each set of multiple pregnancy siblings, of the three-cluster solution profiles using in-model and out-of-model variables**

|  | Variables | Subgroup 1 | Subgroup 2 | Subgroup 3 | p-value |
| --- | --- | --- | --- | --- | --- |
| In-model Variables | **SRS-2** | 18.50 (13.75) | 36.00 (11.75) | 57.00 (25.00) | <0.001 |
|  | **EmQue: Emotion Contagion** | 0.17 (0.43) | 0.17 (0.46) | 0.50 (0.67) | 0.003 |
|  | **EmQue: Attention to others’ emotions** | 1.43 (0.43) | 1.29 (0.57) | 1.29 (0.43) | 0.187 |
|  | **EmQue: Prosocial actions** | 1.33 (0.33) | 1.00 (0.33) | 0.83 (0.50) | <0.001 |
|  | **BRIEF: Emotion control** | 13.00 (5.00) | 14.00 (4.00) | 18.00 (5.00) | <0.001 |
|  | **BRIEF: Inhibit** | 21.00 (5.50) | 25.00 (7.00) | 30.00 (9.00) | <0.001 |
|  | **BRIEF: Shift** | 11.00 (2.00) | 13.00 (2.00) | 17.00 (4.50) | <0.001 |
|  | **BRIEF: Working memory** | 21.00 (4.00) | 26.00 (5.75) | 30.00 (8.50) | <0.001 |
|  | **BRIEF: Plan & organize** | 13.00 (3.00) | 16.50 (3.00) | 17.00 (4.00) | <0.001 |
|  | **GA** | 29.79 (3.68) | 30.50 (3.29) | 29.86 (4.71) | 0.003 |
|  | **IMD**, | 12.88 (15.34) | 23.00 (17.43) | 14.40 (9.62) | 0.018 |
|  | **Days TPN** (0:1:2), n= | 13:39:10 | 15:23:13 | 30:19:9 | 0.003 (X^2^=16.41) |
|  | **Days ventilation** (0:1:2), n = | 34:24:4 | 19:18:14 | 32:22:4 | 0.013 |
|  | **Days CPAP** (0:1:2), n= | 7:39:16 | 2:34:15 | 18:36:4 | 0.001 |
| Out-of-model variables | **SDQ total** | 5.00 (5.00) | 8.17 (5.75) | 13.00 (6.50) | <0.001 |
|  | **SDQ: Internalizing** | 1.00 (2.00) | 2.00 (3.00) | 5.00 (4.00) | <0.001 |
|  | **SDQ: Externalizing** | 3.50 (3.00) | 6.00 (3.75) | 9.00 (5.50) | <0.001 |
|  | **CSPS** | 18.90 (2.00) | 17.84 (4.00) | 17.00 (3.00) | 0.006 |
|  | **WPPSI: Full scale IQ** | 112.00 (16.00) | 105.50 (22.25) | 100.00 (30.00) | 0.007 |
|  | **CBQ: Negative affectivity** | 3.58 (1.15) | 4.25 (0.80) | 4.55 (1.08) | <0.001 |
|  | **CBQ: Surgency** | 5.00 (0.73) | 4.75 (1.25) | 4.58 (1.30) | 0.071 |
|  | **CBQ: Effortful control** | 5.67 (0.83) | 5.17 (0.92) | 5.08 (1.42) | <0.001 |
|  | **Corrected age at assessment: years** | 4.60 (0.50) | 4.69 (0.94) | 4.60 (0.49) | 0.778 |
|  | **IMD at assessment** | 12.02 (15.26) | 18.49 (17.88) | 15.29 (18.66) | 0.203 |
|  | *Sex (M:F), n=* | 32:30 | 24:27 | 30:28 | 0.859 (X^2^=0.30) |
|  | *Total, n (%)* | 62 | 51 | 58 | / |
| *Note*: *Median (IQR) is provided unless otherwise stated. P-values refer to results from Kruskal-Wallis nonparametric* test*.* Days TPN (0:1:2), days ventilation (0:1:2) and days CPAP (0:1:2), correspond to the ratio of the three clinical risk categories: 0,1 and 2. Respectively, they correspond to zero days, more than zero days but less than the top quintile and within the top quintile. ^$^=one missing participants; ^$$^=two missing participants. Chi-squared test was used for categorical comparisons when subject count per cell was >5 and Fisher’s Exact when cell count was 5 or less.  Abbreviations: *BRIEF = Behavior Rating Inventory of Executive Function; CBQ = Childhood Behavioral Questionnaire; CSPS = Cognitively Stimulating Parenting Scale; CPAP = continuous positive airway pressure. EmQue = Empathy Questionnaire; GA = gestational age; IMD = Index of Multiple Deprivation; IQR = interquartile range; PMA = post menstrual age at scan; SDQ = Strengths and Difficulties Questionnaire; SRS-2 = Social Responsiveness Scale – Second Edition; TPN = total parenteral nutrition.* | | | | | |

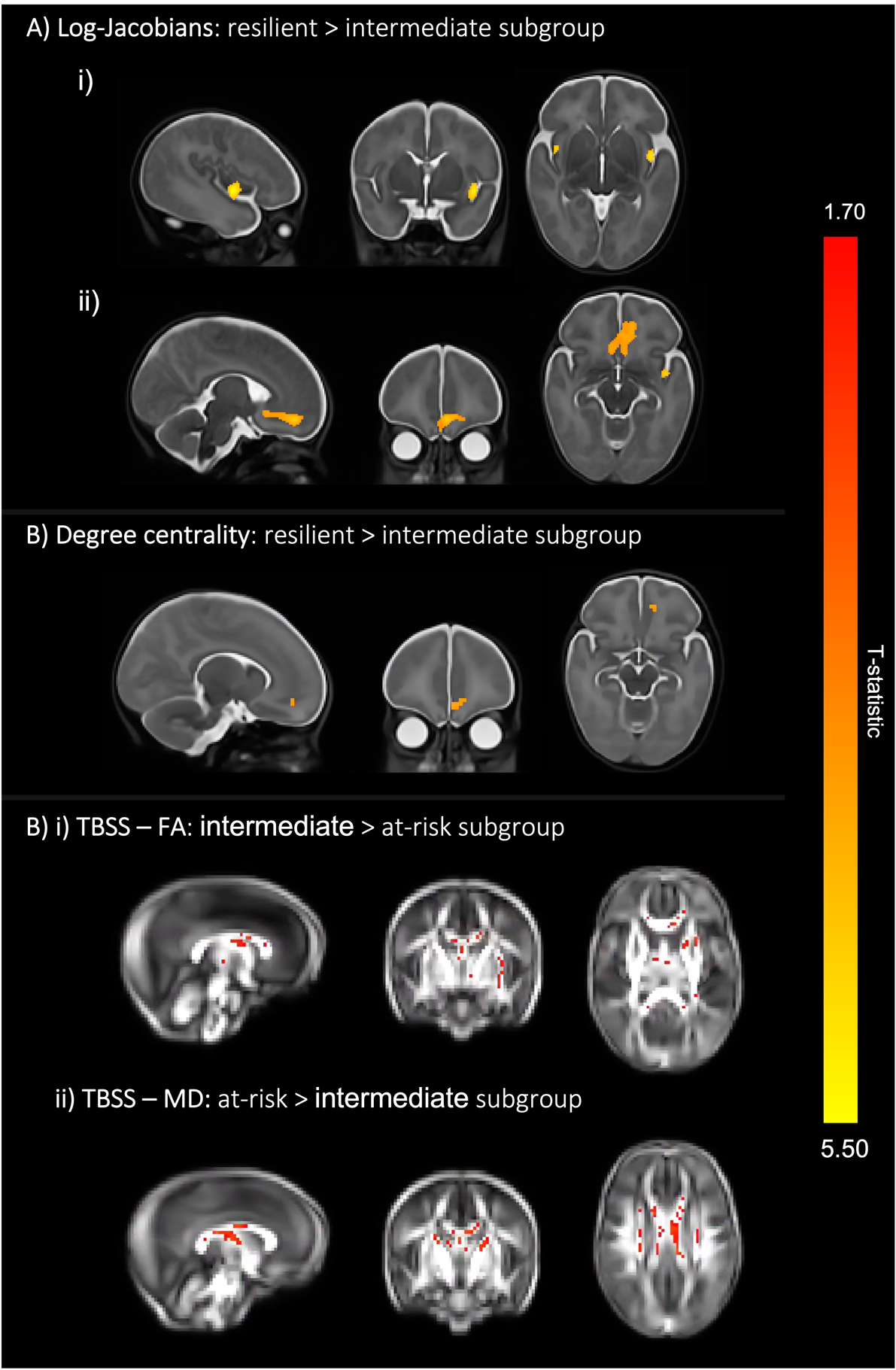

**Figure S6. Brain differences at term-equivalent age of the three cluster-solution, including only one child from each set of multiple pregnancy siblings.** A) Compared to subgroup 3, subgroup 1 had voxels in the i) left and right insula and the ii) left and right orbitofrontal cortex with significantly larger log-Jacobian determinant values at p < .05 (i.e., larger relative volumes in these areas). Sex and PMA were included as covariates and TFCE and FWE corrections were applied. B) an overlapping left orbitofrontal region had voxels showing larger degree centrality (i.e., higher functional connectivity with all other voxels in the grey matter mask) values in subgroup 1 compared to subgroup 3 (*p*=0.08). Sex and PMA were included as covariates and FWE and TFCE corrections were applied. T-statistic values are represented in the color bar, where red colored voxels indicate lower T-statistic values and more yellow voxels indicate higher T-statistic values. C) Colored voxels indicate regions of white matter with: i) significantly higher FA values in the intermediate subgroup compared to the at-risk subgroup and ii) significantly higher MD values in the at-risk subgroup compared to the intermediate subgroup (*p*<0.05). T-statistic values are represented in the color bar, where red colored voxels indicate smaller T-statistic values and yellow voxels indicate higher T-statistic values, ranging between 1.70 and 5.50.

For C=2, out of the 24 sibling sets with more than one sibling included in the clustering analyses, four sibling sets (Figure S7; sibling sets: D, F, P and R) do not cluster together, while the remaining 20 sibling sets do. As for C=3, 5 out of the 24 sibling sets do not co-cluster (Figure S7; sibling sets: D, J, P, S and W). Although siblings tend to group together, we think it is unlikely they may be driving the subgroup profiles described in the main analyses, as we find similar results in the sensitivity analyses which only include one sibling from each set. Moreover, sibling sets are evenly dispersed across the different subgroups and do not cluster into a single subgroup, which suggests that children born from multiple pregnancies do not tend to co-cluster with one another.

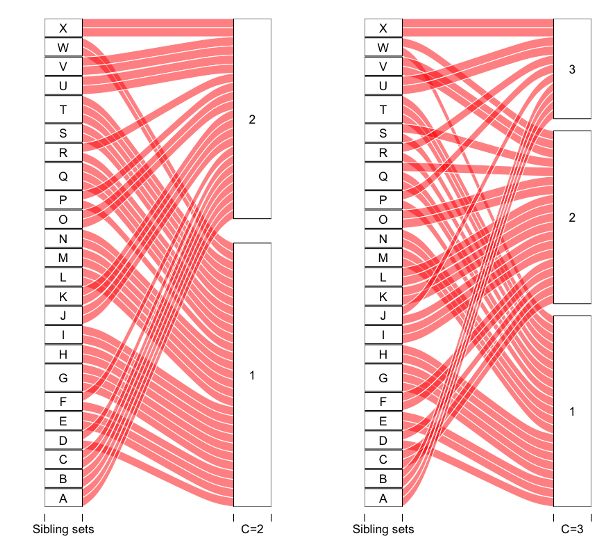

**Figure S7. Alluvial plots for sibling sets.** Alluvial plots indicating which subgroup each sibling within the sibling sets clusters into. Results for the two-cluster solution (C=2) are on the left and the three-cluster solution (C=3) on the right.

**Investigating differences between subgroup profiles after adjusting for confounders**

To explore whether possible confounders (age and sex) altered our results we re-ran comparisons between clusters for non-MRI out-of-model (SDQ internalizing , SDQ externalizing, CSPS, full scale IQ, CBQ surgency, CBQ effortful control and CBQ negative affectivity) and in-model measures (SRS-2 total, EmQue emotion contagion, EmQue Attention to others’ emotions, EmQue Prosocial actions, BRIEF Emotion control, BRIEF Inhibit, BRIEF Shift, BRIEF Working memory, BRIEF Plan and organize). We found that results remained similar and survived FDR correction. However, CBQ surgency scores were lower in the at-risk compared to the resilient subgroups in the C=2 cluster solution only, after adjusting for age and sex. This was also the case for C=3, whereby the at-risk subgroup showed significantly lower CBQ surgency scores compared to both resilient and intermediate subgroups. A meta-analysis investigating gender differences in temperament found that boys show increased surgency levels compared to girls (^26^). Furthermore, differences between the at-risk and intermediate subgroups in EmQue prosocial subscale scores emerged after adjusting for age and sex. All results survived FDR correction.
